# Local nucleosome dynamics and eviction following a double-strand break are reversible by NHEJ-mediated repair in the absence of DNA replication

**DOI:** 10.1101/866673

**Authors:** Vinay Tripuraneni, Gonen Memisoglu, Wei Zhu, Trung Tran, Alexander J Hartemink, James E Haber, David M MacAlpine

**Affiliations:** Department of Pharmacology and Cancer Biology, Duke University Medical Center, Durham, NC 27710; Department of Biology and Rosenstiel Basic Medical Sciences Research Center, Brandeis University, Waltham, MA 02454; Department of Computer Science, Duke University, Durham, NC 27708

## Abstract

Although the molecular events required for the repair of double-strand breaks (DSB) have been well characterized, the role of epigenetic processes in the recognition and repair of DSBs has only been investigated at low resolution. We rapidly and synchronously induced a site-specific DSB in *Saccharomyces cerevisiae* upstream of the *PHO5* locus, which has well-positioned nucleosomes. Utilizing MNase-seq epigenome mapping we interrogated the order of chromatin changes that occur immediately following a DSB by generating a base-pair resolution map of the chromatin landscape. In wild type cells, the first nucleosome left of the break was rapidly evicted. The eviction of this flanking nucleosome was dynamic and proceeded through an early intermediate chromatin structure where the nucleosome was repositioned in the adjacent linker DNA. Other nucleosomes bordering both sides of the break were also shifted away from the break; however, their loss was more gradual. These local changes preceded a broader loss of chromatin organization and nucleosome eviction that was marked by increased MNase sensitivity in the regions ∼8 kb on each side of the break. While the broad loss of chromatin organization was dependent on the end-processing complex, Mre11-Rad50-Xrs2 (MRX), the early remodeling and repositioning of the nucleosome adjacent to the break was independent of the MRX and YKU70/80 complexes. We also examined the temporal dynamics of NHEJ-mediated repair in a G1-arrested population, where 5’ to 3’ end-resection of DSB ends is blocked. Concomitant with DSB repair, we observed the re-deposition and precise re-positioning of nucleosomes at the originally-occupied positions. This re-establishment of the pre-lesion chromatin landscape suggests that a DNA replication-independent mechanism exists in G1 cells to preserve epigenome organization following DSB repair.

## Introduction

Failure to repair DNA double strand breaks (DSB) results in cell cycle arrest and ultimately programmed cell death, while improper repair can lead to profound alterations or loss of genomic information through translocations, inversions, deletions and other genomic aberrations (Hanahan and Weinberg 2000; Halazonetis et al. 2008). Accurate and timely recognition and repair of DSBs is critical to maintaining genomic integrity, and must be accomplished within the context of chromatin (Hauer and Gasser 2017). The canonical access-repair-restore model (Soria et al. 2012; Polo and Almouzni 2015) for DNA repair in eukaryotic genomes recognizes that chromatin presents a significant obstacle for DNA double strand break recognition and repair processes. Chromatin must thus undergo obligatory remodeling to evict existing nucleosomes and allow break-recognition factors to access and signal for repair complexes to assemble and mend the broken DNA. While repair of the broken DNA ensures that genome integrity is preserved, chromatin architecture must also be restored to preserve the integrity of the epigenome.

Much of our understanding of DNA DSBs in the context of chromatin structure has been informed by experiments at the yeast *MAT* locus (Haber 2012). This locus is specifically and efficiently cut by the HO endonuclease *in vivo*, and has served as a model system for many genetic and molecular investigations of DNA damage and repair (Sugawara and Haber 2006; Haber 2012). Immediately following a DSB, the MRX complex recognizes the broken ends of DNA and recruits the downstream kinase Tel1 which, in conjunction with Mec1, phosphorylates histone H2A (analogous to H2AX in mammalian cells) at up to 50 kb on either side of the break (Shroff et al. 2004; Kim et al. 2007; Lee et al. 2014). Moreover, nucleosomes surrounding the break site are evicted in an Mre11p-Rad50p-Xrs2p (MRX)-dependent manner up to 8 kb surrounding the break and this process serves to facilitate the access and activity of subsequent repair factors (Tsukuda et al. 2005; Shim et al. 2007). This nucleosome eviction that occurs surrounding a DSB appears to be a conserved phenomenon as similar results are also observed at specific breaks induced in the genomes of mammalian systems (Tsukuda et al. 2005; Berkovich et al. 2007; Goldstein et al. 2013). The MRX complex and Ku70/Ku80 (KU) each complex associate with the broken ends of chromatin both prior to and during the nucleosome remodeling/eviction steps and this association is critical to their roles in the repair of the break via NHEJ (Boulton and Jackson 1996; Moore and Haber 1996). While it is clear that the KU complex retards 5’-3’ end resection and that MRX is critical in the recognition and processing of the end of the DSB (Lee et al. 1998; Symington 2014; Cannavo and Cejka 2014), these roles do not describe the function of MRX or KU complexes relative to nucleosome dynamics and histone octamer eviction immediately following a break.

The two primary pathways to repair DSBs are homologous recombination (HR) and nonhomologous end joining (NHEJ). HR involves the generation of single-stranded DNA (ssDNA) around the DSB via end-resection machinery and subsequently requires the presence of a homologous sequence of DNA (most commonly a sister chromatid) to serve as a template that can be used to resynthesize DNA sequences. However, except in special circumstances, such as *MAT* switching where the donor is heterochromatic, with highly positioned nucleosomes that are not found at the recipient locus (Weiss and Simpson 1998; Ravindra et al. 1999), it is difficult to distinguish the chromatin structures of homologous regions undergoing recombination (Hicks et al. 2011; Tsabar et al. 2016). NHEJ, however, is a simpler process and requires no template for repair of the DNA strand. Instead, the Yku70-Yku80 heterodimer along with MRX recognizes both ends of a DSB and then in conjunction with Lif1 and Nej1, the enzyme Dnl4 catalyzes their re-ligation (Boulton and Jackson 1996; Wilson et al. 1997; Teo and Jackson 1997; Lieber 2010). Though the process of reconstituting the DNA backbone is straightforward at the molecular level, it remains unclear how the local chromatin environment is perturbed and restored following a break and how similar this chromatin architecture is to the pre-lesion state.

In both yeast and higher eukaryotic systems, it has been demonstrated that specific chromatin assembly factors and chromatin remodelers are associated with and required for proper repair and restitution of the chromatin state following a DSB (Chai and Huang 2005; Tsukuda et al. 2005; Shim et al. 2007; Liang et al. 2007; Kim and Haber 2009; Neumann et al. 2012; Horigome et al. 2014; Kwon et al. 2015; Polo 2015). However, chromatin remodelers may exhibit pleiotropic activities such as regulation of cell-cycle dependent gene expression and may not directly influence the chromatin environment surrounding a DSB. Thus, despite a critical role for ATP-dependent chromatin remodeling complexes in DSB repair, the precise structural outcomes of nucleosome eviction and subsequent reassembly and positioning of these nucleosomes to their final pre-lesion state has yet to be elucidated. Further it has been challenging to discriminate between replication-independent chromatin reestablishment and the replication-coupled chromatin reassembly that occurs following S-phase (Tsabar et al. 2016). While DNA replication-dependent mechanisms of chromatin reassembly following break repair are expected to restore the original parental chromatin organization (Li and Tyler 2016), the fidelity and accuracy of replication-independent mechanisms for histone octamer deposition have not been fully explored. Specifically, it is unknown if an independent mechanism is competent to restore the complex regulatory landscape, including transcription factor binding and nucleosome positioning, to its pre-damage state.

We sought to interrogate, at nucleotide-resolution, the spatiotemporal kinetics of chromatin remodeling and eviction in response to a single DSB and the subsequent restoration of chromatin organization following replication-independent NHEJ mediated repair. We engineered an inducible site-specific DSB system in *S. cerevisiae* by inserting the 117 bp HO endonuclease recognition site 579 bp upstream of the *PHO5* locus (**Figure 1A**). The *PHO5* gene in yeast is marked by a very well positioned +1 nucleosome that is responsible for regulating its expression and has been frequently used as a model for well-defined chromatin organization (Almer and Hörz 1986; Almer et al. 1986; Schmid et al. 1992; Gaudreau et al. 1997; Hertel et al. 2005; Tsabar et al. 2016). Upon break induction, we found that local nucleosome eviction and repositioning within the first 200 bp around the DSB at *PHO5* occurs prior to the broad pattern of nucleosome eviction associated with a persistent DSB. We also observed that the eviction of a single nucleosome immediately adjacent to the DSB was not dependent on *MRE11* suggesting that while local end binding of MRX at a DSB is rapid (Zhang et al. 2007), this event is neither sufficient nor necessary to induce the early chromatin changes at a DSB. Following NHEJ-mediated repair of this break near *PHO5*, we observed that the broad pattern of chromatin accessibility was similar to unbroken DNA and that local chromatin architecture was restored rapidly. This replication-independent reassembly yields positioning of nucleosomes around the DSB that recapitulates the chromatin state prior to DSB induction suggesting that an active and structurally accurate mechanism exists to preserve chromatin architecture following DNA damage and repair.

**Figure 1.**
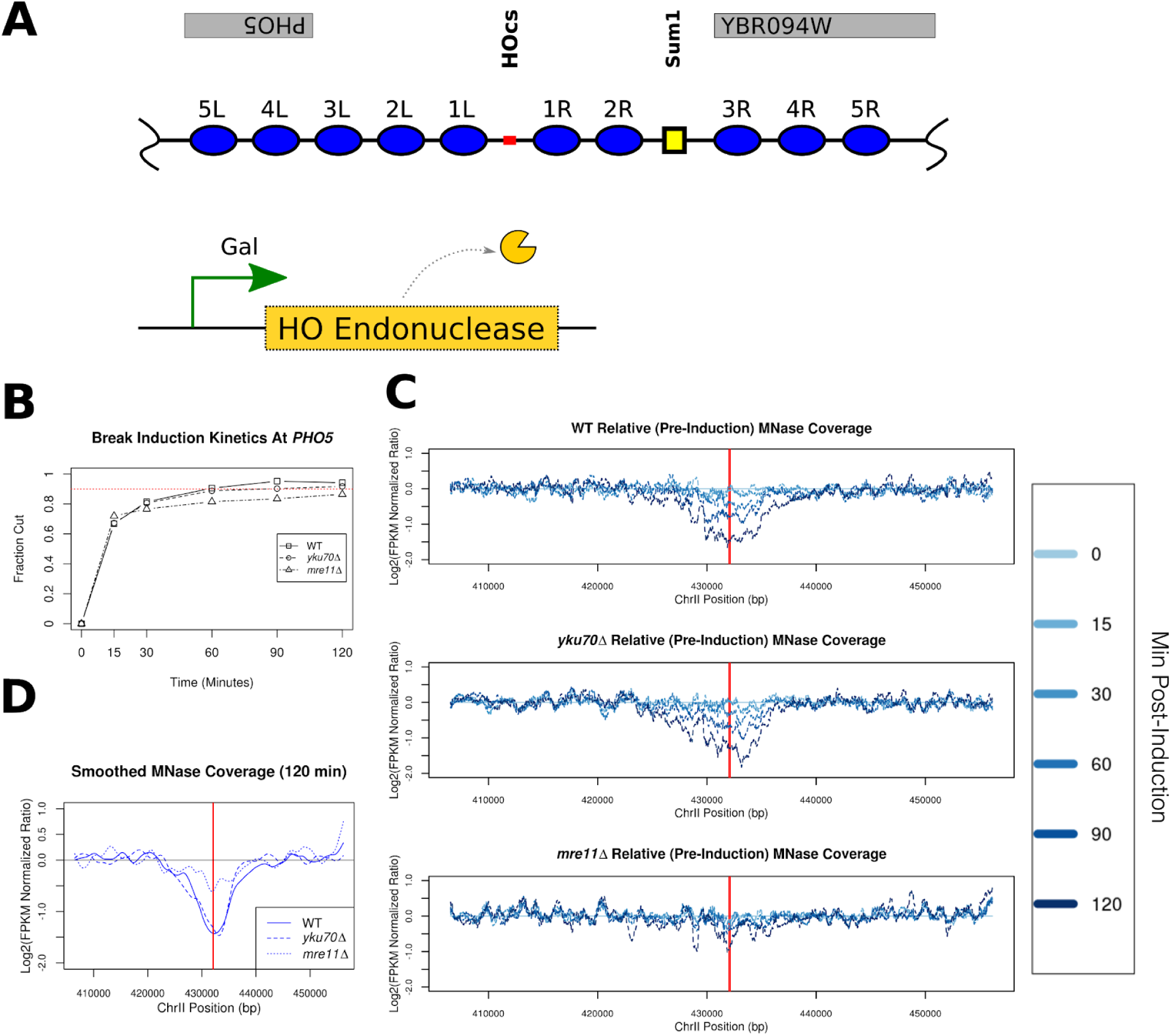
Inducible DSB break at PHO5. **A**. Schematic illustrating the ectopic 117bp HO cut site(HOcs) that was engineered 579 bp upstream of the endogenous *PHO5* locus on Chr II (marked in red). The +1 nucleosome of *PHO5* is annotated 4L in this illustration. A predicted Sum1 protein binding site is illustrated with a small yellow square. Expression of the HO endonuclease is regulated by an inducible promoter. Nucleosomes are numbered and annotated based on their distance and orientation (left/right) relative to the HO cut site. **B**. Southern blot analysis of HO endonuclease induced cutting near *PHO5* with fraction of cut DNA (y-axis) plotted as a function of time (x-axis) for the WT, *yku70Δ* and *mre11Δ* strains. Dotted red line depicts cleavage of 90% of the DNA. **C.** Chromatin accessibility surrounding the break greatly increases for WT, *yku70Δ*, but is limited in *mre11Δ* strains. Greater time post-induction (min) is plotted in darker shades of blue indicated in the legend (right). **D.** Direct comparison between the WT and two mutant strains for the terminal (120 min) MNase sensitivity (in **C**) which has been smoothed.

## Results

### Site-specific inducible DSB at the PHO5 locus

The eviction of histone octamers from the broken chromatin and subsequent remodeling of nucleosomes are required steps in the recognition and processing of double strand breaks (Polo and Almouzni 2015; Hauer and Gasser 2017). To understand the relationship between the spatial and temporal kinetics of the chromatin remodeling events that accompany a DSB, we inserted a 117bp HO endonuclease recognition sequence upstream of the *PHO5* locus (**Figure 1A**), which is marked by a well positioned +1 nucleosome (nucleosome 4L in **Figure 1A**) and has been frequently used as a model for well-defined chromatin organization (Almer and Hörz 1986; Almer et al. 1986; Schmid et al. 1992; Gaudreau et al. 1997; Hertel et al. 2005; Tsabar et al. 2016). Insertion of the HO recognition site did not impact the positioning of the +1 nucleosome at *PHO5*, but did have a modest effect on local chromatin structure **(Supplemental Figure 1**). We noticed that nucleosomes flanking the HO recognition site were better positioned than in the native locus. Using a galactose-inducible HO endonuclease, we find that the *PHO5* 117bp HO sequence cuts just as efficiently as the endogenous *MAT* locus (Tsukuda et al. 2005; Lee et al. 1998) with 70% of the target locus being cut within 15 min and >90% within 1 hour of galactose induction as measured by southern blot quantification (**Figure 1B & Supplemental Figure 2).** We also observe similar cutting kinetics in strains lacking *MRE11* or *YKU70*. If HO expression is turned off, NHEJ can readily repair DSB ends; but when HO is continually expressed nearly all DSBs persist (Moore and Haber 1996; Aylon et al. 2004).

In response to a persistent DSB, histone octamers are broadly evicted from the DNA at a rate of ∼4kb/h, symmetrically from the DSB (Tsukuda et al. 2005). The DSB is then subject to NHEJ-mediated repair or 5’ to 3’end-resection for subsequent homology-driven repair (White and Haber 1990; Tran et al. 2004; Tsukuda et al. 2005; Zhu et al. 2008; Mimitou and Symington 2008; Symington and Gautier 2011; Jasin and Rothstein 2013). We first profiled the temporal changes in chromatin organization surrounding the DSB using sensitivity to micrococcal nuclease (MNase) as a proxy for chromatin accessibility. We found a time-dependent increase in chromatin accessibility (loss of sequence fragment coverage) following induction of the break in WT cells (**Figure 1C**; top panel). A similar increase in chromatin accessibility surrounding the break was observed in the absence of *YKU70 (***Figure 1C**; middle panel); however, loss of *MRE11 (***Figure 1C**; bottom panel) abrogated the increased chromatin accessibility as has been previously reported at lower resolution (Tsukuda et al. 2005). By 120 min, chromatin accessibility surrounding the break had increased nearly 4 fold relative to pre-induction, and spread ∼16 kb in WT (with ∼8 KB on either side of the break) and ∼20 kb in *yku70Δ* cells (with ∼10 KB on either side of the break). We observed only a minimal degree of increased accessibility (less than 2 fold relative to pre-induction) at the break site in *mre11Δ* cells which spreads less than 10kb (∼ 5KB in either direction from the break) **(Figure 1D**). These results establish our *PHO5 GAL::HO* inducible system as a robust model to interrogate chromatin surrounding DSBs.

### MNase-seq epigenome mapping resolves chromatin dynamics at base-pair resolution

We sought to precisely define the spatiotemporal dynamics of octamer eviction and chromatin remodeling at nucleotide resolution following induction of a DSB at the *PHO5* locus. In order to assess changes in chromatin occupancy, we generated genome-wide chromatin occupancy profiles (GCOPs) (Henikoff et al. 2011; Belsky et al. 2015; Gutiérrez et al. 2018), by coupling paired-end sequencing with MNase digestion to reveal the factor-agnostic location of both nucleosomes and smaller DNA binding factors. DNA protected by the histone octamer will yield sequenced fragments of ∼150 bp; whereas smaller DNA binding proteins such as transcription factors will protect fragments between 20 and 80 bp. GCOPs are visualized by plotting the fragment length as a function of chromosomal coordinate for the midpoint of each recovered fragment (**Figure 2A**); thus, well-positioned and phased nucleosomes appear as tight and evenly arrayed clouds of data points with a fragment size of ∼150 bp. Similarly, DNA protected by smaller sequence-specific DNA binding factors are represented by a focus of smaller sized fragments. We chose an unrelated locus on chromosome IV containing two genes, *CTH1* and *GIR2*, with well-defined chromatin organization, to serve as an internal control representing static chromatin for these studies. This locus contains nine well positioned nucleosomes and a clearly defined Aft2 footprint upstream of the *CTH1* gene body (**Figure 2B**) that do not change over the time course. In contrast, the *GAL7* locus exhibits a dramatic change in chromatin organization in response to the addition of galactose (**Figure 2C**). These changes include the rapid eviction and dephasing of the nucleosomes within the gene body due to high levels of transcription (Yarger and Hopper 1979; Lohr et al. 1995; Platt and Reece 1998). To further quantify our results, we employed a two-dimensional cross correlation analysis with an idealized nucleosome (**Supplemental Figure 3**, & detailed in methods) to define the position, occupancy and fuzziness (correlation score with an idealized nucleosome; lower scores being more fuzzy) for each nucleosome. These metrics are summarized in pictograph form for both the *CTH1/GIR2* and *GAL7* loci (**Figure 2D & E).** Specifically, the location, size and color opacity of the circles represent position, occupancy and fuzziness. The occupancy, positioning and fuzziness of the nucleosomes at *CTH1/GIR2* remain stable and unchanged throughout the time course (**Figure 2D**), while we observe profound chromatin architectural changes at *GAL7* following induction (**Figure 2E**). These findings are consistent across all strains profiled in this study (**Supplemental Figure 4 & 5**) and demonstrate the specificity and sensitivity of GCOPs for describing the spatiotemporal dynamics of chromatin occupancy.

**Figure 2.**
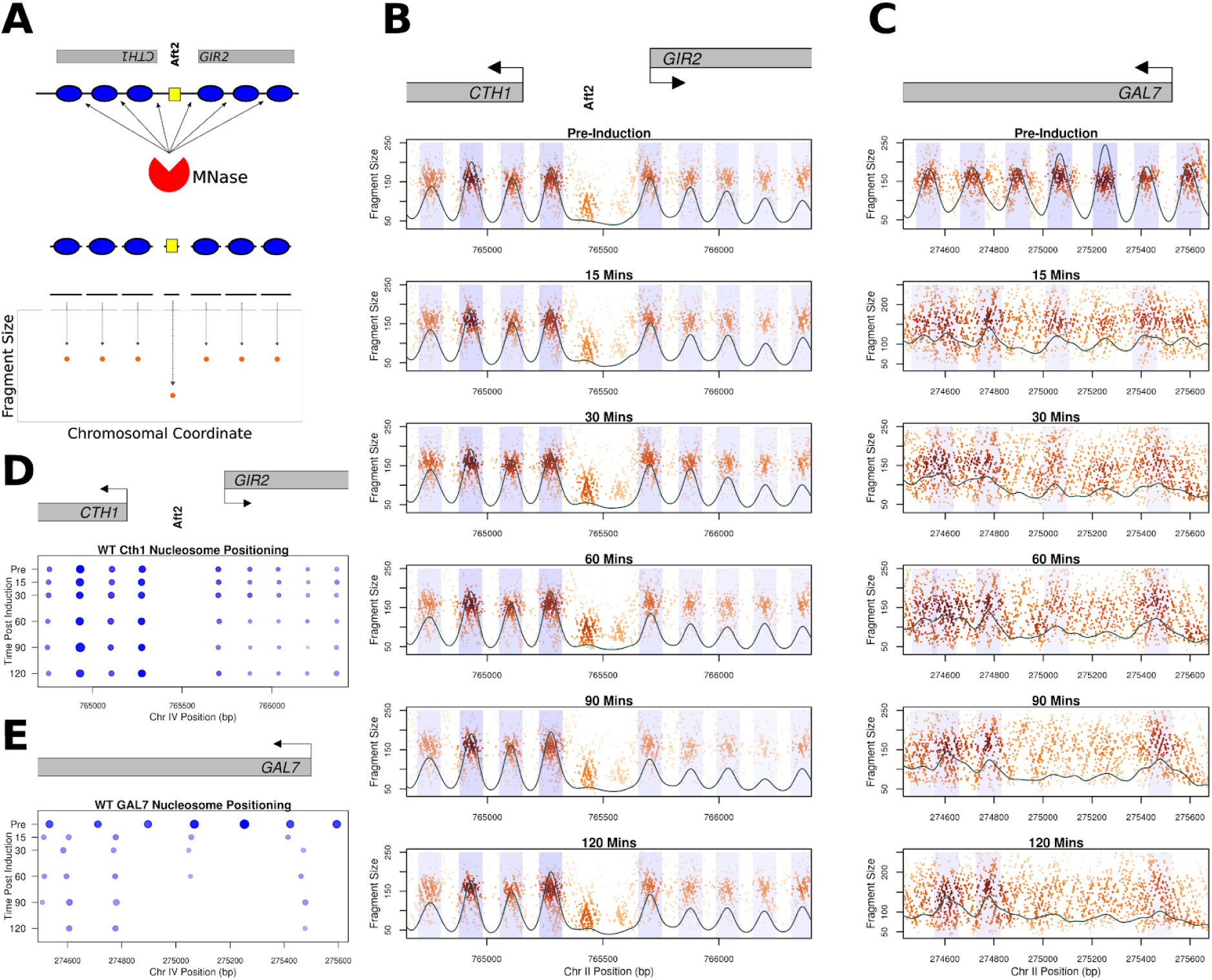
Genome-wide Chromatin Occupancy Profiles (GCOPs) to detect chromatin changes at base pair resolution. **A.** Schematic of the MNase-Seq based assay to generate GCOPs. MNase is used to digest unprotected DNA, and the subsequent nucleosome and TF protected fragments of DNA are subjected to paired-end sequencing and the fragment size is plotted as a function of the position of its midpoint along the chromosome. **B.** GCOP for the *CTH1* and *GIR2* loci which serve as a control for unaltered chromatin. Gray boxes depict gene bodies, with arrows indicating the direction of transcription. The bold vertical text annotates the Aft2p binding site in between these two genes. A two-dimensional cross correlation with an idealized nucleosome (see methods) is calculated at every base pair and depicted as a continuous black trace. The peaks of this cross-correlation analysis represent the most likely position of a locally detected nucleosome and we have shaded +/− 0.5 standard deviation of nucleosome position from the center of each detected peak. The intensity of the shaded color is proportional to the nucleosome’s peak cross-correlation score. **C.** GCOP for the *GAL7* locus following galactose induction. **D** and **E** serve as pictographic summaries of the chromatin changes observed where each blue circle represents a corresponding nucleosome from the plots in **B and C**. The size of a given circle is proportional to its occupancy; its x position is the location of the peak of the 2D density kernel cross correlation score and the intensity of color is proportional to the peak cross-correlation value (fuzziness).

### High-resolution analysis of DSB-induced chromatin changes at PHO5

To interrogate chromatin structure in response to a DSB at the *PHO5* locus, GCOPs were generated for wild-type, *yku70Δ* and *mre11Δ* mutant strains **(Figure 3).** Prior to DSB induction (**Figure 3;** pre-induction; top panels), all strains exhibit a similar chromatin organization, with 5 nucleosomes positioned both left and right of the HO cut site (HOcs) (illustrated in **Figure 1A**). In wild-type cells we observe the rapid loss of the 1L nucleosome upon galactose induction. The 1R and 2R nucleosomes shift away from the break and towards each other, and are positioned so closely together after 30 min that they are difficult to discern as individual nucleosomes. The crowding of the 1R and 2R nucleosomes appears to be limited by a small DNA factor binding site (presumably Sum1) since the 3 nucleosomes downstream of this footprint do not alter their positioning following break induction in wild-type or mutant strains. We also observe a time-dependent loss of reads within the HO cut site as well as an increase in smaller length fragments on either side of the cut site. This small factor footprint is not symmetric however; it is larger on the left side of the break and resides at the sequence previously occupied by the evicted 1L nucleosome (**Figure 3A**, blue arrowhead **& Supplemental Figure 6**). The presence of this footprint is also seen in both *yku70Δ* and *mre11Δ* mutants suggesting that is not dependent on these DSB end binding factors (**Supplemental Figure 6**). Additionally, we observe a gain in nucleosomal-sized fragments in the linker region between the 1L and 2L nucleosomes (denoted by the red arrowhead) suggesting the sliding of histone octamers away from the break. **(Figure 3**).

**Figure 3:**
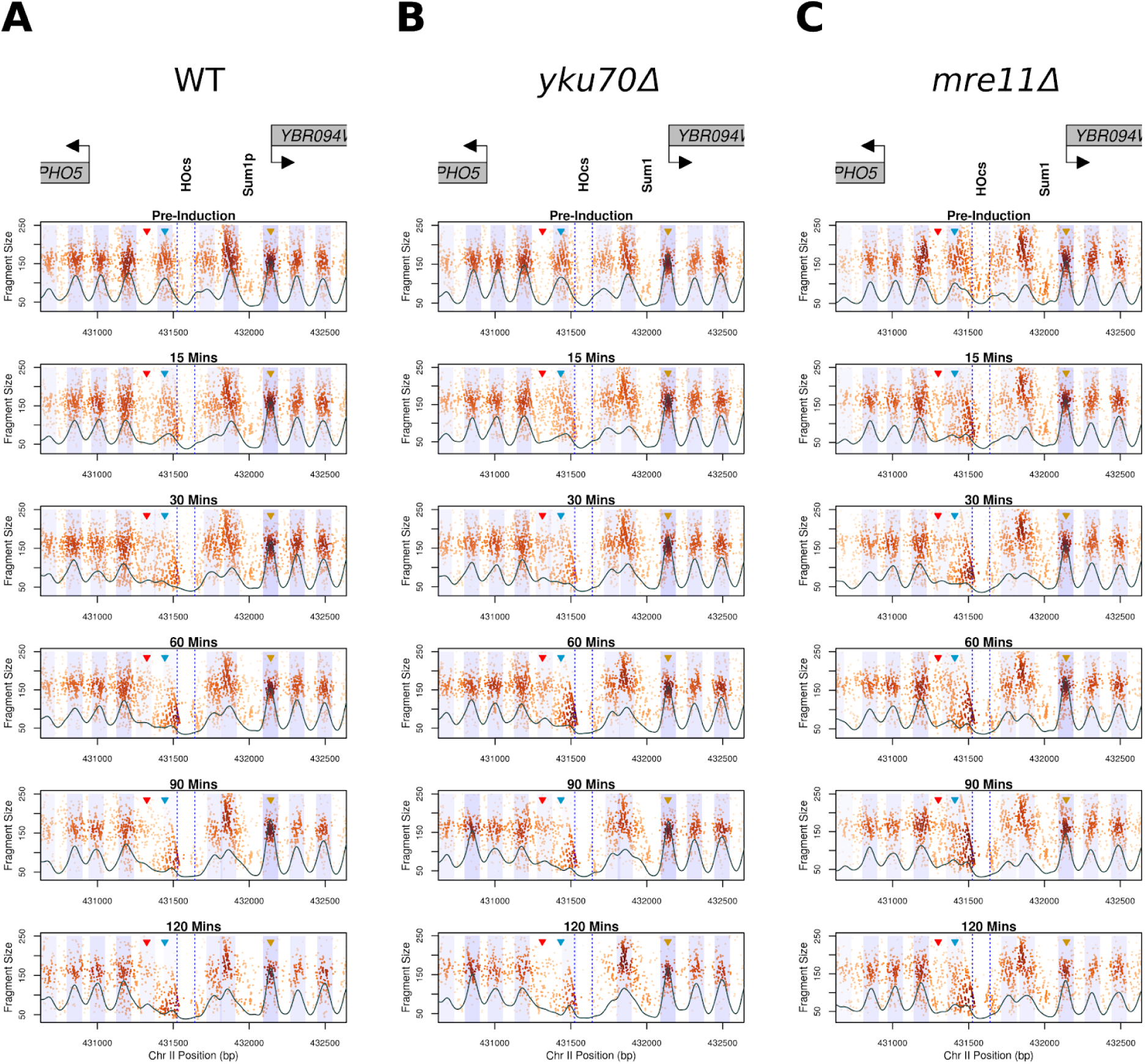
Local chromatin changes in response to a DSB at the *PH05* locus. GCOPs of WT (**A**) *yku70Δ* (**B**) and *mre11Δ* (**C**) strains over the experimental time course. Gene bodies are annotated above the plots in gray, with arrows indicating the direction of transcription. Dotted blue lines denote the 117 bp HO recognition site and a predicted Sum1 protein binding site is also depicted. Similar to Figure 2B **& C**, a 2D kernel cross-correlation score is depicted as a continuous black line. Peaks of the cross correlation score were called and the corresponding area in the plot is shaded +/− 0.5 standard deviations of nucleosome width in light blue. The transparency of this shaded region is proportional to the peak value of the cross correlation score with larger (stronger) values being relatively intensely shaded and smaller (weaker) values being relatively lightly colored. The regions quantified in Figure 4 are marked by colored arrowheads for the 1L (blue) and 3R (gold) nucleosomes the 1L-2L linker (red) region.

To summarize our observations in **Figure 3**, we generated pictograms for nucleosome positioning, occupancy and fuzziness over the experimental time course (**Figure 4A**). The pictogram captures the rapid eviction of the 1L nucleosome, a leftward shift for nucleosomes 2L, 3L and 4L as well as the rightward shift and crowding of nucleosomes 1R and 2R. **Figure 4A**). Over the course of 120 min following induction, we observe a time-dependent decrease in the overall occupancy. Despite this reduction in nucleosome occupancy, nucleosome positioning, does not change appreciably when compared to early post-induction time points (<30 min), indicating that the nucleosomes which remain following induction of the break retain their early post-break positioning (**Figure 4A**).

**Figure 4.**
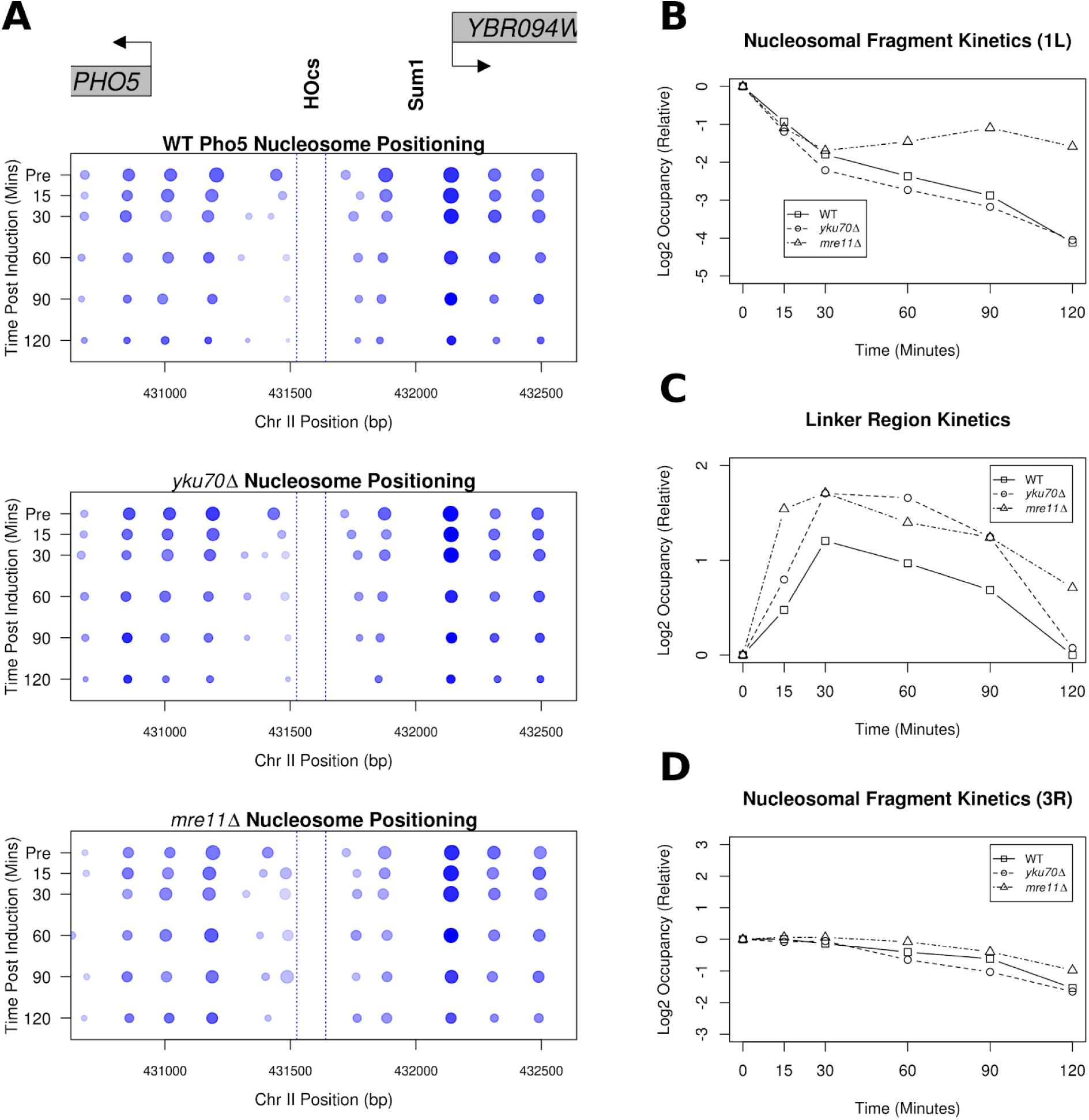
*MRE11* is important for local nucleosome dynamics at a DSB. **A.** Pictographs depicting the chromatin changes near *PHO5* following DSB induction. Increasing time is plotted downward on the y axis and the boundaries of the 117-bp HO recognition site are denoted by dotted blue lines. **B.** The FPKM corrected relative occupancy of the 1L nucleosome (Figure 3; left blue arrowhead) for the WT, *yku70Δ* and *mre11Δ* mutant strains. **C.** Occupancy of the 1L-2L linker region was quantified as in B (Figure 3; red arrowhead). **D.** The same relative occupancy analysis performed in **B** is plotted for the 3R nucleosome (Figure 3; green arrowhead). This nucleosome is positioned immediately to the right of a predicted Sum1 protein binding site (Figure 1A).

Using 2D density kernel cross-correlation analysis, we are unable to algorithmically detect the 1L nucleosome later than 15 min following break induction in wild-type and mutant strains (**Figure 3**). However, this cross-correlation metric assumes a homogeneous single-state system (i.e. an idealized nucleosome precisely localized in the bulk of the population) and it fails to detect more complex state mixtures (i.e. partial nucleosomes, poorly positioned nucleosomes, occupancy by either a nucleosome or a transcription factor, etc); thus, significant changes in nucleosome occupancy between wild-type and mutant strains may become obscured by low 2D cross correlation values. Given the apparent competition between the 1L nucleosome being evicted, the accumulation of smaller protected fragments over the same sequence, as well as the gain in nucleosomal sized fragments in the linker region between the 1L and 2L nucleosomes, we sought to employ a more sensitive and direct method to analyze the data. We directly inferred the kinetics of 1L nucleosome eviction illustrated in **Figure 3** by measuring fragment loss at the pre-induction 1L nucleosome position **(Figure 4B**) and similarly assessed the occupancy of the 1L-2L nucleosomal linker region by measuring fragment gain at this position (**Figure 4C**). As a local control we also analyzed the 3R nucleosomal fragments **(Figure 4D).** These metrics demonstrated that the nucleosomes immediately around the break were dynamic; whereas the 3R nucleosome behaved remarkably consistently across wild-type and mutant strains. Although the loss of the 1L nucleosome is similar in all strains up to 30 min, at later time points (60 min and beyond) we detected delayed 1L nucleosome loss in the *mre11Δ* strain relative to the loss observed in the wild-type and *yku70Δ* strains (**Figure 4B**). Interestingly, the kinetics of nucleosome appearance in the 1L-2L linker region--either by histone deposition or nucleosome sliding--were consistent between wild-type and the mutant strains (**Figure 4C**). The relative fragment loss observed at the 1L nucleosome position parallels the relative gain in nucleosome fragments in the 1L-2L linker region in all strains, suggesting that this 1L nucleosome is initially shifted left of the break. Together, these results indicate that the 1L nucleosome and the 1L-2L linker region experience early occupancy changes independent of the MRX and Yku70-Yku80 binding proteins, and that these alterations precede the eventual nucleosome loss and broad changes in chromatin accessibility we observe at later time points.

### Genome-wide changes in response to a single DSB

Despite the early perturbations in nucleosome positioning that were localized to the immediate vicinity of the break, the broad in increase in chromatin accessibility surrounding the break at terminal time points led us to question if any other distant chromatin changes had occurred. Given that our data provides nucleosome occupancy and position throughout the entire genome at base-pair resolution, we sought to interrogate the post-induction nucleosome occupancy changes genome-wide in response to a single, persistent DSB. We determined the relative ratio between the 2D cross correlation score (similarity to an idealized nucleosome) for genic nucleosomes in the pre-induction state versus the 120 min post-induction state and discovered that the most significant changes were confined to the genes within ∼8kb the DSB and the *GAL* loci on chromosome II **(Supplemental Figure 7**). The subtle changes in nucleosome occupancy that were detected on the remaining fifteen unbroken chromosomes, were likely associated with perturbations in gene expression elicited by a DSB (Lee et al. 2000). In contrast to the changes in nucleosome occupancy that spread 8 kb from the break, changes in nucleosome positioning were constrained to the sequences immediately surrounding the break (**Supplemental Figure 8**). Taken together these findings suggest that the structure of the chromatin landscape (except for the region immediately around the DSB) is largely preserved across the genome in response to a DSB.

### Replication independent restoration of chromatin following NHEJ

How is the local eviction and nucleosome displacement restored following repair of a DSB? Specifically, how rapidly is the chromatin structure restored and does it return to the pre-induction state? It remains unclear whether chromatin organization is precisely restored in a replication-dependent or independent manner following NEHJ (Chiruvella et al. 2013; Geuting et al. 2013; Emerson and Bertuch 2016; Gao et al. 2016; Li and Tyler 2016). To profile the changes that occur following repair via NHEJ, we arrested the cells in the G1 phase of the cell cycle by the addition of α-factor for 3.5 h and then induced HO expression for 1 h with galactose to ensure complete cutting and chromatin disruption at the target locus. Repair and potential chromatin reassembly at *PHO5* was allowed to occur by adding 2% dextrose to repress *GAL::HO* expression (**Figure 5A & Supplemental Figure 10**). In the absence of continued transcription, the HO endonuclease is rapidly degraded, allowing NHEJ to re-join the 4-nt 3’ overhanging ends (Moore and Haber 1996; Kaplun et al. 2000). In G1-arrested cells, 5’ to 3’ resection is retarded in WT cells and replication is blocked (Ira et al. 2004; Aylon et al. 2004; Zierhut and Diffley 2008; Clerici et al. 2008). This condition allowed us to specifically interrogate replication-independent chromatin reassembly following NHEJ repair. Following repression of *GAL::HO*, we observed a progressive increase in the fraction of DNA being repaired, with rejoining of approximately 50% occurring within 6 h. We confirmed that this repair was *DNL4-*dependent and thus represented classical NHEJ (Teo and Jackson 1997; Wilson et al. 1997) (**Figure 5A & Supplemental Figure 9).**

**Figure 5.**
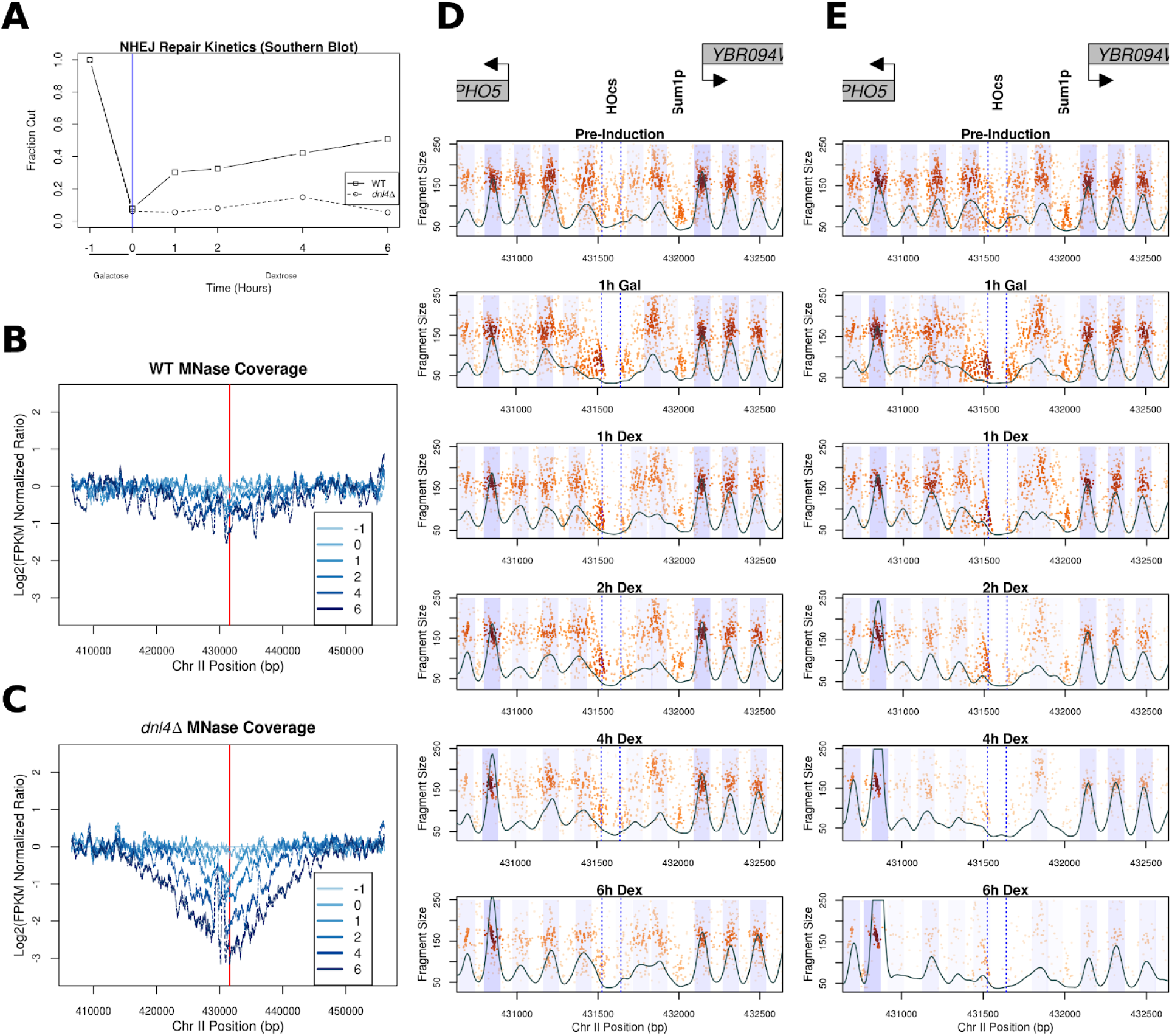
NHEJ mediated DSB repair of the *PHO5* region prevents broad nucleosome eviction. **A**. Cutting/rejoining kinetics the previously described ectopic 117-bp HO recognition site that was engineered upstream of the *PHO5* gene. Suppression of HO (starts at the time denoted by the blue line) facilitates repair by NHEJ, a Dnl4-dependent repair process, in this donorless system. **B & C**. Bulk MNase coverage over the cut/repair time course in the WT strain (**B**) and the *dnl4Δ* mutant (**C**). The red line denotes the HO cut site. **D & E.** Chromatin occupancy profiles of the WT (**D**) and *dnl4Δ* (**E**) strains. As in Figure 3, gene bodies are annotated above the plots in gray and the 2d kernel cross-correlation score is plotted in black over this region. Time pre-induction (−1), after HO induction (0), and after repression of HO expression (1, 2, etc.) are shown. The boundaries of the HO cut site are denoted by dotted blue lines. Peaks of the cross correlation score were called and the corresponding area in the plot is shaded +/-0.5 standard deviation of nucleosome width in light blue.

We first examined broad chromatin accessibility via sensitivity to MNase digestion throughout the process of NHEJ and noted only a modest two-fold increase in chromatin accessibility at the sequences surrounding the break. These changes were consistent with the reconstitution of chromatin on the 50% of remaining DNA that underwent repair (**Figure 5B**). In contrast, we observed substantially more chromatin accessibility and loss of DNA surrounding the break in a *dnl4Δ* strain (**Figure 5C**), consistent with the broken DNA ends being slowly degraded in the absence of a competent repair pathway (Aylon et al. 2004). Importantly, we were able to verify transcriptional repression of *GAL::HO* by generating GCOPs for the *HO* gene body where we observed the restoration and maintenance of pre-induction nucleosome structure within the HO gene body following dextrose addition (**Supplemental Figure 10**). Lastly, we confirmed the G1 arrest of the experimental cell population for the duration of the experiment via FACS (**Supplemental Figure 11**).

We also examined the spatiotemporal dynamics of chromatin structure in high resolution at the sequences immediately surrounding the DSB in both wild-type and the *dnl4Δ* strain. The chromatin structure was similar in the pre-induction state and following HO induced cutting for 1 h. Following repression of *HO*, we begin to detect NHEJ repair within 1 h and the reappearance of the 1L nucleosome in the wildtype strain (**Figure 5D**); this was not observed in the *dnl4Δ* strain suggesting that repair of the DNA was necessary and sufficient for restoring the 1L nucleosome (**Figure 5 E**). We also observe that the 1R and 2R nucleosomes assume their pre-induction positioning, and the overall chromatin state at 6 h post-dextrose recapitulates that of the pre-induction state in the wildtype strain (**Figure 6A**; top panel) while in the *dnl4Δ* strain we observed a profound loss of recovered fragments suggesting much of the DNA surrounding the broken locus has been degraded. Despite the extensive histone eviction and resectioning that occurred in the *dnl4Δ* strain by the terminal time point, at earlier time points (up to 2 h post dextrose addition), the chromatin organization and structure around the break was similar to the chromatin structure in the wild type strain. This indicates that the break-induced changes are stable until degradation or repair of the broken chromosome and not dependent on the continued expression of HO endonuclease (**Figure 5E and 6A;** bottom panel). We again quantified the occupancy kinetics of the 1L and 3R nucleosomes in these G1-arrested cells and we observe a close association between the loss of the 1L nucleosome and near complete cutting of the *PHO5* locus (**Figure 6B**). The eviction and re-deposition of the 1L nucleosome closely parallels the break and subsequent repair of the DNA in wildtype strains, but is never restored in the *dnl4Δ* strain (**Figure 6B**). In contrast, the 3R nucleosome exhibits a decrease in occupancy reflective of the proportion of DNA that remains unrepaired by NHEJ (**Figure 6B**). Furthermore, the kinetics of reappearance or deposition of the 1L nucleosome along the chromatin parallels the fraction of DNA repaired (**Figure 5A & Figure 5D**).. This suggests that the processes intrinsic to NHEJ-mediated DSB repair are competent to facilitate the reestablishment and positioning of nucleosomes to their pre-lesion arrangement along the chromatin in a replication-independent but repair-coupled manner (**Figure 6C**).

**Figure 6.**
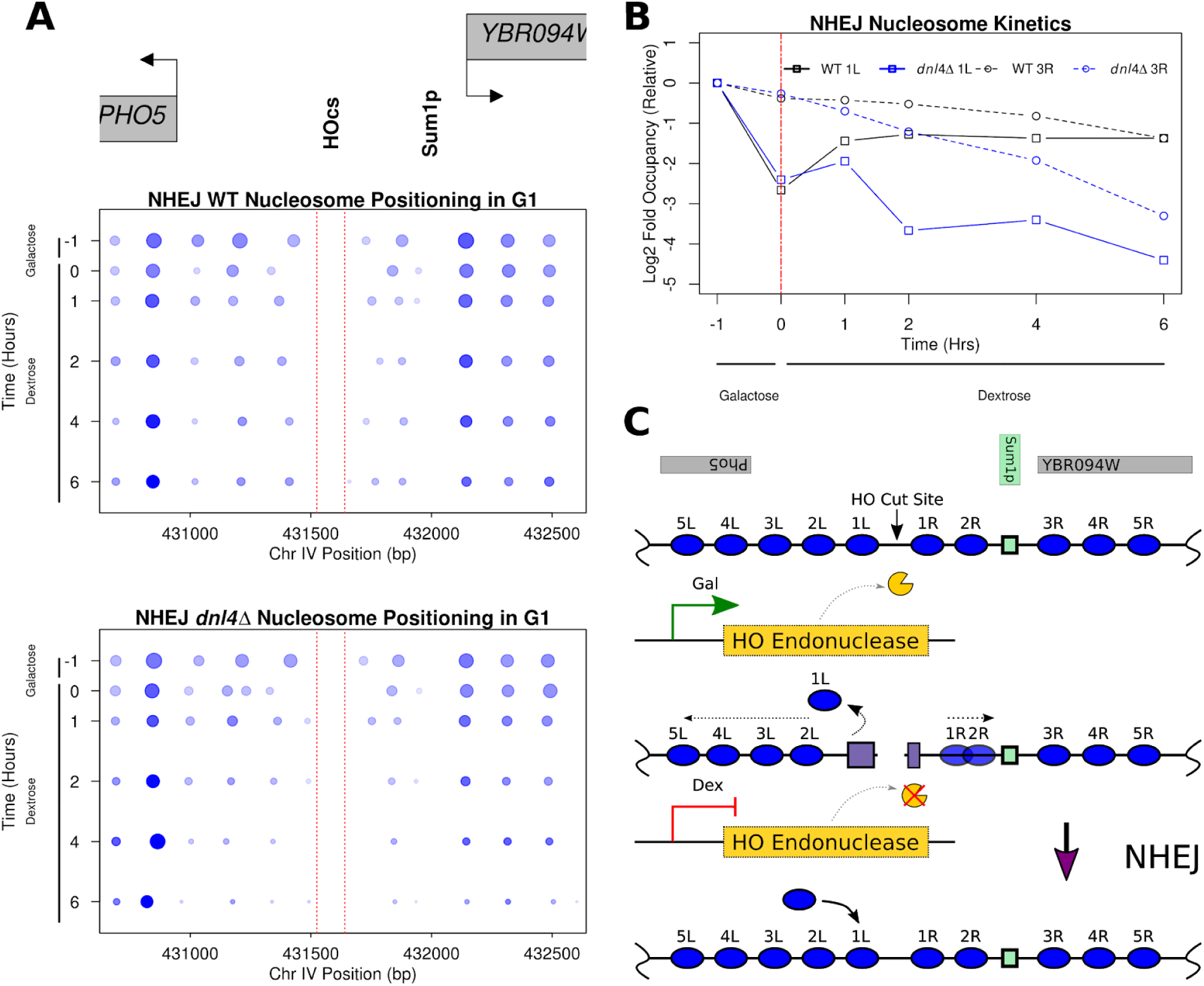
NHEJ mediated DSB repair restores chromatin structure independent of replication. **A**. Pictographs depicting local nucleosome changes over the experimental time course in the WT and *dnl4Δ* strains. Similar to the analysis performed in Figure 4 we precisely quantified the alterations in chromatin structure following break induction and NHEJ mediated repair. Again, increasing time is plotted downward on the y axis and the boundaries of the HO cut site are denoted by dotted blue lines. **B.** The log2-relative (to pre-induction) occupancy of the 1L (squares/solid line) and 3R (circles/dotted line) nucleosomes (denoted 1L and 3R respectively in Figure 6C) are plotted over time for this experiment in black (WT) and blue (*dnl4Δ*). **C.** Model for the replication-independent nucleosome reassembly and chromatin restoration following NHEJ.

## Discussion

The structural changes that chromatin undergoes following a double-strand break and its ensuing repair have only been investigated at low-resolution (Shim et al. 2005; Tsukuda et al. 2005; Tsabar et al. 2015). To interrogate the kinetics of chromatin dynamics following a DSB and throughout NHEJ mediated repair we developed a system that would allow us to survey the genome-wide chromatin occupancy changes at nucleotide resolution with an inducible site-specific break upstream of the *PHO5* locus. We identified an early cascade of discrete chromatin changes that occurred immediately following break induction which were independent of *MRE11*. Specifically, the rapid eviction of the 1L nucleosome which may have proceeded through an intermediate remodeling event that saw displacement of the 1L nucleosome into the adjacent linker region. These immediate changes in chromatin structure at the break were followed by the *MRE11*-dependent broader eviction of histone octamers. The early and local changes to chromatin were reversible through repair of the genetic lesion by NHEJ, and occurred in a replication-independent manner.

Previous investigations of chromatin structure surrounding a DSB have relied on ChIP or PCR/probe based methods to query the presence of a specific sequences/loci (Tsukuda et al. 2005; Shim et al. 2007; Goldstein et al. 2013; Tsabar et al. 2016; Li and Tyler 2016). These methods have established a low-resolution model of broad nucleosome eviction dependent on MRX activity and facilitated by Sgs1, Exo1, Dna2, Fun30, and Ino80 (Dubrana et al. 2007; Mimitou and Symington 2008; Eapen et al. 2012; Westmoreland and Resnick 2015). This combination of chromatin remodeling complexes, helicases and nucleases suggests that these broad chromatin changes are related to and coupled with DNA end-resection (Symington and Gautier 2011). At 120 min post-induction, we observe approximately a two-fold reduction in the occupancy of nucleosomes to the left and right of the DSB in WT and YKU70-strains but this occupancy loss is abrogated in the MRE11-strain. This degree and extent of MNase sensitivity we observed at our terminal time point (120 min)--which in wild-type strains extends approximately ∼8 kb from the break--is consistent with prior reports (Tsukuda et al. 2005). During this eviction and resection phenomena, we do not observe any shifts or positioning changes in the remaining nucleosomes despite their loss in occupancy. Similarly, when we examined the remaining fifteen unbroken chromosomes we were also unable to detect any significant changes in nucleosome occupancy or positioning.

These data suggest that the structure of the chromatin landscape may be largely preserved up until the moment of eviction and resection. While changes in global transcript levels, elevated rates of histone turnover, higher order changes in nuclear organization, and changes in chromatin fiber condensation following double-strand breaks have been described (Lee et al. 2000; Hauer and Gasser 2017; Amitai et al. 2017; Hauer et al. 2017; Seeber and Gasser 2017), our work suggests that the fundamental organization of chromatin (at the level of the individual nucleosome) does not undergo radical remodeling and re-positioning immediately following a break. Preservation of pre-existing chromatin structure surrounding the break and throughout the genome may be critical for maintaining regulated gene expression and preventing potential spurious transcription.

Prior to these broad and late changes to chromatin structure, rapid (within 15 – 30 min) local alterations to nucleosome positioning and occupancy were observed concomitant with break induction. Specifically, we documented the repositioning of a single nucleosome to the left of the break, the gain of small-factor footprints immediately bordering both sides of the break, and the local dephasing of nucleosomes around the break. The ATP-dependent chromatin remodeling complexes INO80, SWR1 and RSC have been implicated in facilitating repair/recognition factor access to DSB ends(Morrison et al. 2004; Tsukuda et al. 2005; Shim et al. 2007; Liang et al. 2007; Horigome et al. 2014). However, the immediate and early changes (<60 min) were independent of *MRE11* and implicate another biochemical mechanism or chromatin remodeling process in the early phase of a DSB or, alternately, the chromatin changes at the ends of the break are entropically driven.

In concert with these early local chromatin changes, we were able to detect the appearance of a small factor footprint on both sides of the DSB within 15 mins following induction of the DSB. We initially hypothesized that these small factor occupancy footprints were the result of canonically associated DSB recognition and repair factors. The MRX complex and YKU70-80 heterodimer are known to rapidly (∼20 min) associate with the ends of broken DNA (Palmbos et al. 2008; Wu et al. 2008). However, this footprint’s asymmetry and appearance was independent of *MRE11* or *YKU70*, suggesting that this footprint is the result of another DNA end binding factor. We investigated the sequences bordering the DSB and found no well-described motifs that would explain the observed footprint or account for differential digestion of the recovered up/downstream sequence following MNase digestion. The creation of two new free DNA ends at a DSB serves as a substrate for chromatin remodeling complexes such as RSC and INO80--which have previously described to be associated with DSBs with rapid kinetics (Liang et al. 2007; Shim et al. 2007). While the recruitment of INO80 is thought to be MRX-dependent, the RSC complex may interact with the broken DNA to create space for recognition and repair factors by sliding nucleosomes away from the DSB and also prevent histone octamers from sliding towards the free ends and off the DNA. The displacement of the H2A/H2B dimer immediately around a DSB may also suggest the presence H3-H4 tetrasomes at a DSB and potentially explain the smaller sized footprint we observe (Shroff et al. 2004). Finally, we also considered the possibility that that the HO endonuclease itself associates with the ends of the 117bp HO recognition sequence following break induction, and HO itself could be responsible for this footprint; however, the asymmetric small-factor footprint remains long after dextrose repression and degradation of HO in a *dnl4Δ* strain (Kaplun et al. 2000).

Following recognition and repair of a DSB, the concomitant re-establishment of chromatin structure surrounding the repaired locus is critical for maintaining epigenome integrity. Previous studies with HO-induced DSBs have demonstrated the re-establishment of chromatin structure following recombinatorial repair of the break; a process which necessitates template driven new DNA synthesis and requires specific chromatin remodeling machinery and histone chaperones (Tsabar et al. 2016; Mehta et al. 2017). The process of NHEJ mediated repair, however, intrinsically lacks a DNA synthesis-coupled chromatin assembly step and it is unclear if the pre-lesion chromatin structure and organization is capable of being re-established prior to the next S-phase. Chromatin remodeling complexes--such as HIRA, RSC-- and histone chaperones--such as CAF-1 and ASF-1— have been implicated in NHEJ mediated repair and in the re-establishment of H3 histone occupancy at the repaired break (Shim et al. 2005; Linger and Tyler 2005; Kim and Haber 2009; Li and Tyler 2016). These studies suggest an active mechanism to reestablish chromatin occupancy following NHEJ mediated DSB repair, but the structure of this repaired chromatin specifically at the level of individual nucleosome occupancy and positioning has yet to be elucidated. Furthermore, the implication of a replication associated histone chaperone, specifically CAF-1, in re-establishing histone occupancy following NHEJ mediated repair in cycling cells suggests that this process might yet be coupled to a DNA synthesis or necessitate a replication-dependent re-assembly of the chromatin. The notion of a replication-dependent reset of chromatin following repair of a DSB is augmented by work which has demonstrated that the dephosphorylation of H2A--a DSB coupled modification which extends up to 50kb away from the DSB--is most efficient on free histone dimers and not in the DNA packaged nucleosome (Keogh et al. 2006; Nakada et al. 2008). Taken together these observations postulate a model that that re-establishment or resetting of the chromatin state following repair of a DSB by NHEJ is replication-coupled.

By arresting donorless haploid cells in G1 with ɑ-factor throughout the entire experiment we were able to prevent replication and eliminate the possibility of repair via sister chromatid recombination. The break-induced eviction, dephasing and small-factor footprints that we previously described as a consequence of a DSB are all reversible within 1 hour of NHEJ mediated repair and are most clearly evident at 6 hours post repair, when a majority of the broken DNA has been re-ligated. This suggests that repair-coupled but replication-independent chromatin machinery is competent to reset chromatin structure following a DSB and occurs at a rate limited by repair of the genetic lesion. In the broader context of DNA repair, particularly in higher eukaryotes, it is important to consider that NHEJ (and related/derivative non-template mediated end-joining processes) are the predominant and rapid repair pathways used to mend a DSB (Mao et al. 2008). Given that cells experiencing genotoxic stress, and DSBs in particular, may be post-mitotic, our work suggests that a concerted and rapid chromatin reassembly mechanism exists coupled to NHEJ which serves to maintain the structure of the epigenome in the absence of DNA replication or synthesis.

## Materials and Methods

### Strains

We built our model system in strain JKM139 lacking *HML*, *HMR* (Sandell and Zakian 1993; Lee et al. 1998). By selecting survivors on a YEP-GAL plate we recovered alterations of the *MAT***a** cleavage site (designated HOcs deleted) that could no longer by cut by the endonuclease (Lee et al. 1998). Strains used in this study have the following genotypes:

| Strain | Genotype |
| --- | --- |
| DM579 | *hoΔ hmlΔ::ADE1 MATa* (HOcs Deleted)<br>*hmrΔ::ADE1 ade1-100 leu2-3,112*<br>*trp::hisG lys5 ura3-52 ade3::GAL::HO*<br>*bar1Δ::ADE3 ChrII:431525::HOcs(117bp)* |
| DM581 | *hoΔ hmlΔ::ADE1 MATa* (HOcs Deleted)<br>*hmrΔ::ADE1 ade1-100 leu2-3,112*<br>*trp::hisG lys5 ura3-52 ade3::GAL::HO*<br>*bar1Δ::ADE3 ChrII:431525::HOcs(117bp)*<br>*dnl4Δ::TRP1* |
| DM645 | *hoΔ hmlΔ::ADE1 MATa* (HOcs Deleted)<br>*hmrΔ::ADE1 ade1-100 leu2-3,112*<br>*trp::hisG lys5 ura3-52 ade3::GAL::HO*<br>*bar1Δ::ADE3 ChrII:431525::HOcs(117bp)*<br>*yku70Δ::HPH* |
| DM622 | *hoΔ hmlΔ::ADE1 MATa* (HOcs Deleted)<br>*hmrΔ::ADE1 ade1-100 leu2-3,112*<br>*trp::hisG lys5 ura3-52 ade3::GAL::HO*<br>*bar1Δ::ADE3 ChrII:431525::HOcs(117bp)*<br>*mre11Δ::TRP1* |

Single gene deletions were carried out using single-step-PCR-mediated transformation of yeast colonies (Sikorski and Hieter 1989). Markerless insertion of the 117bp MAT**a** HO cut site (HOcs) (Kostriken and Heffron 1984) was accomplished through the use of CRISPR/Cas9 locus editing, by using the plasmid pGM031, which contains Pho5-gRNA-f (DM1806) and Pho5-gRNA-r (DM1807) cloned into the BplI cut site in bRA90 plasmid (Anand et al. 2017). Primer and gBlock sequences are provided in **Supplemental Table 1**.

### Galactose Inductions

Cells were grown to an OD600 of approximately 0.4-0.5 in YEP with 2% raffinose and 0.1% dextrose. 20% galactose was added to a final concentration of 2% in the medium to induce HO expression. Two 100 ml cell aliquots were harvested at specified time intervals for subsequent southern blotting and chromatin preparation.

### Southern Blotting

Southern blots were performed with DNA probes labeled with αP32-ATP (Perkin Elmer) as previously described (Southern 2006) using Church & Gilbert hybridization buffer (Church and Gilbert 1984). To assess the fraction of DNA cut in the WT, *yku70Δ*, and *mre11Δ* strains, we performed band densitometry analysis (with ImageJ) of each lane and computed the fraction of cut DNA relative to the sum of the cut and uncut DNA bands (**Supplemental Figure 10**). To interrogate the fraction of DNA rejoined via NHEJ, we computed the fraction of each post-induction uncut band to the pre-induction sample (−1 lane in **Supplemental Figure 10**) and then normalized this value to an unrelated control fragment which was probed on Chr IX.

### Cell Cycle Arrest and HO repression

Wild-type cells were grown to an OD600 of approximately 0.2-0.3 in YEP with 2% raffinose and 0.1% dextrose at which point alpha factor was added at a final concentration of 50 ng/ml for 3.5-4 h (GenWay). 20% galactose was added to a final concentration of 2% in the medium to induce HO expression. To repress HO, cells were pelleted at 2000 RPM and then washed twice in YPD + 2% dextrose + 50 ng/ml alpha factor followed by resuspension in YPD + 2% dextrose + 50 ng/ml alpha factor. Two 100 ml cell aliquots were harvested at specified time intervals for subsequent southern blotting and chromatin preparation.

### Flow Cytometry

Flow cytometry was performed on 1mL culture aliquots as previously described (Gutiérrez et al. 2018).

### Chromatin Preparation

Chromatin preparation was performed on cell pellets derived from 100 mL cell aliquots as previously described (Henikoff et al. 2011; Belsky et al. 2015).

### Sequencing Library Preparation

Sequencing libraries were prepared as previously described (Belsky et al. 2015) with the following modifications: NEBNext multiplex oligos for Illumina kit (New England Biolabs) were used in adapter ligation and PCR steps. 12 cycles of PCR were performed as indicated by the NEBNext multiplex oligos for Illumina kit (New England Biolabs). PCR reactions were cleaned using Agencourt AMPure XP beads (Beckman).

### Data Analysis

All figures and plots were generated using R version 3.2.0 and Inkscape version 0.92.2.

### Alignment

Sequencing reads were aligned with Bowtie (Langmead et al. 2009) in paired-end mode to the sacCer3/R64 version of the *S. cerevisiae* and an edited version of Chr II derived from this same genome with the 117 bp HO cut site inserted upstream of *PHO5*.

### Analysis of Chromatin Structure and Occupancy

Sequencing reads from biological replicates for each experiment were merged for all data analysis and sampled to equivalent read depth across each chromosome. Chromatin accessibility was determined for each sample by calculating the total number of reads in a 500bp window stepping every 10 bases −25 kb from the start of the HO recognition site to +25kb of the end of the HO recognition site. The log2-ratio of each sample relative to the pre-induction sample was then calculated to interrogate MNase sensitivity of the region. The boundaries of MNase sensitivity were determined by smoothing the MNase coverage curves and determining the widths of the sensitivity for all strains.

Nucleosomes were called based on a sliding 2-dimensional cross-correlation score at every base to an “idealized” nucleosome 2D density kernel. This kernel was derived by analyzing 8,632 unique nucleosome positions on Chr IV that were mapped utilizing a highly sensitive chemical mapping methodology (Brogaard et al. 2012). This provided us with the approximate size and distribution of reads in our data that corresponded to a canonical well-positioned nucleosome. The variance of the size of the (y) and position of these reads (x) is how we derived the idealized nucleosome 2D kernel. We derived idealized nucleosome kernels for each time-course’s pre-induction sample to control for slight MNase digestion variability across WT and mutant strains. Density plots of these idealized kernels are provided in **Supplemental Figure 3**, and demonstrate the consistency between data sets. Nucleosomes were defined as peaks of this 2D cross-correlation trace above the 10th percentile of all cross-correlation values on Chr IV. We then shaded each nucleosomal peak +/− 0.5 standard deviations of the idealized nucleosome’s x variance to highlight the area of interest. The intensity of this shading is proportional to the maximal nucleosome occupancy within a plotting window time course. Nucleosome occupancy is defined as the number of fragments that fall between 0.5 standard deviations of position (x) and 1 standard deviation of size (y). Nucleosome “fuzziness” is separate from the occupancy and positioning metrics and is defined as the value of the peak of the 2D cross-correlation score. This metric captures how similar or dissimilar a called nucleosome peak is to an idealized nucleosome.

## Data Availability

FASTQ files are deposited at the NCBI SRA (SRA######)

## Acknowledgements

We thank members of the MacAlpine and Haber laboratories for critical comments and suggestions. We also thank Yulong Li for advice on data analysis. This work was supported by the following grants from the National Institute of General Medical Sciences: R35 GM127062 (D.M.M.), R01 GM61766 (J.E.H), R35 GM127029 (J.E.H), R01 GM118551 (A.J.H).

## Supplemental Data and Figures

### Nucleotide sequences

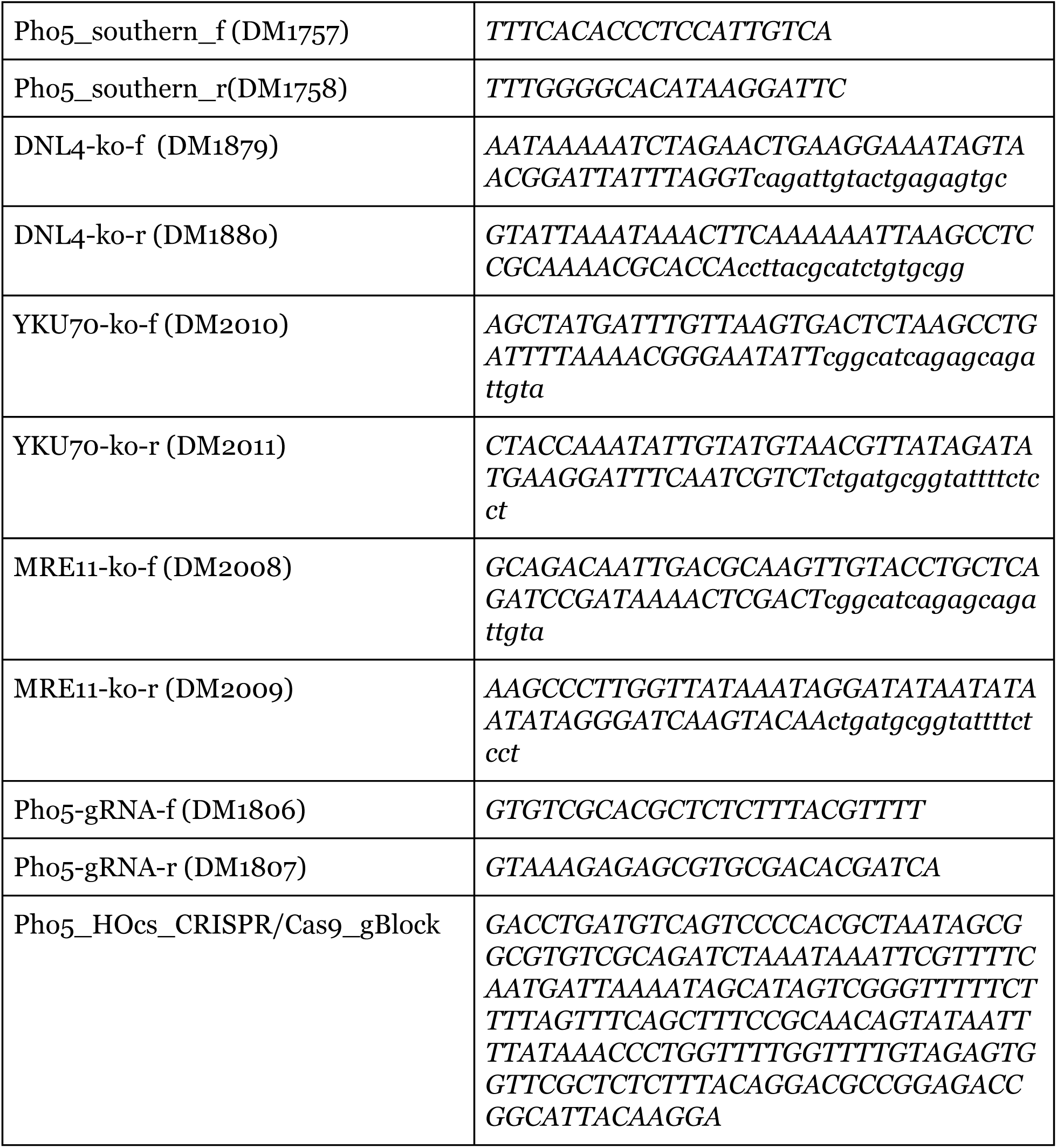

**Supplemental Figure 1.**
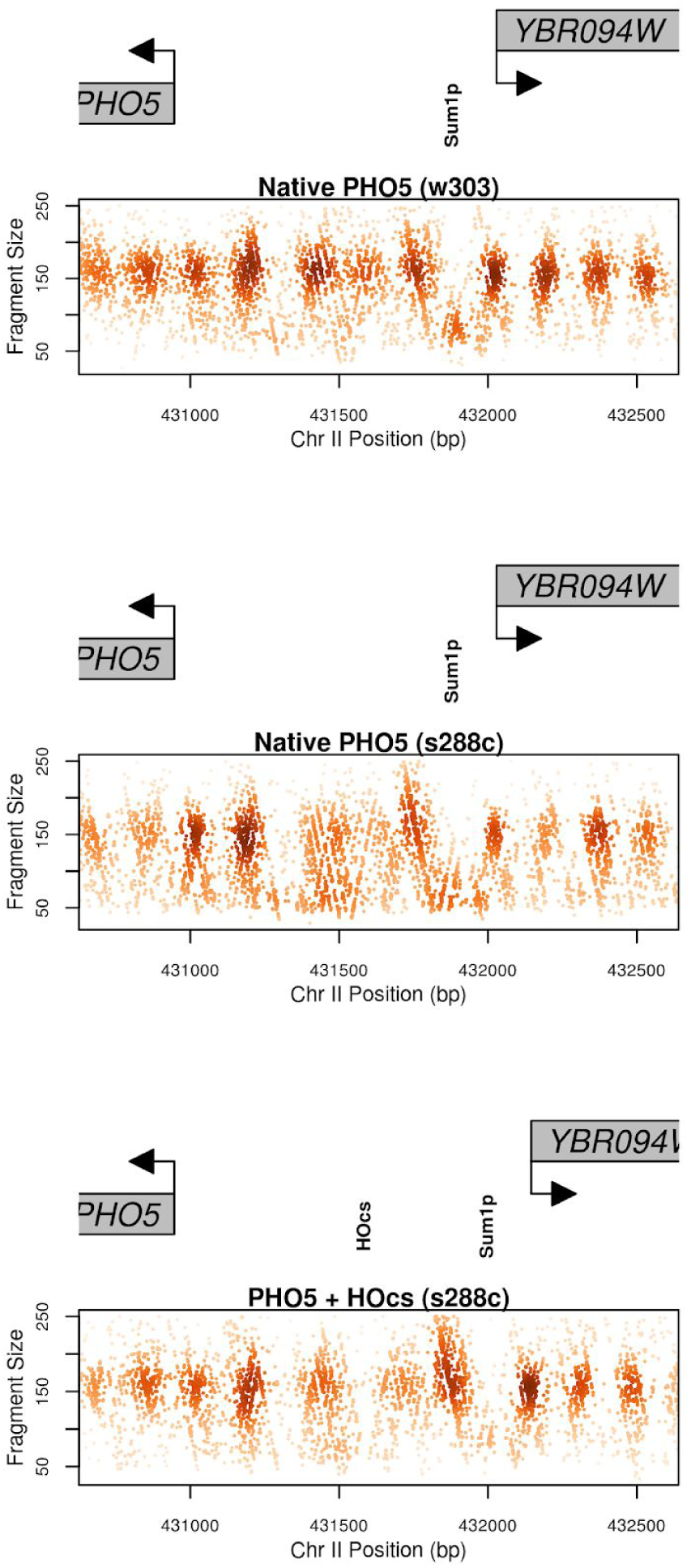
Structure of the *PHO5* locus prior to and after the HOcs insertion. Top panel is the native *PHO5* locus from the w303 yeast strain background while the middle panel is the same locus in the JKM139 strain, which is largely derived from s288c. The bottom panel is the s288c strain with the HOcs insertion. The +1 nucleosome of *PHO5* is the 4L nucleosome in **Figure 1A**.

**Supplemental Figure 2.**
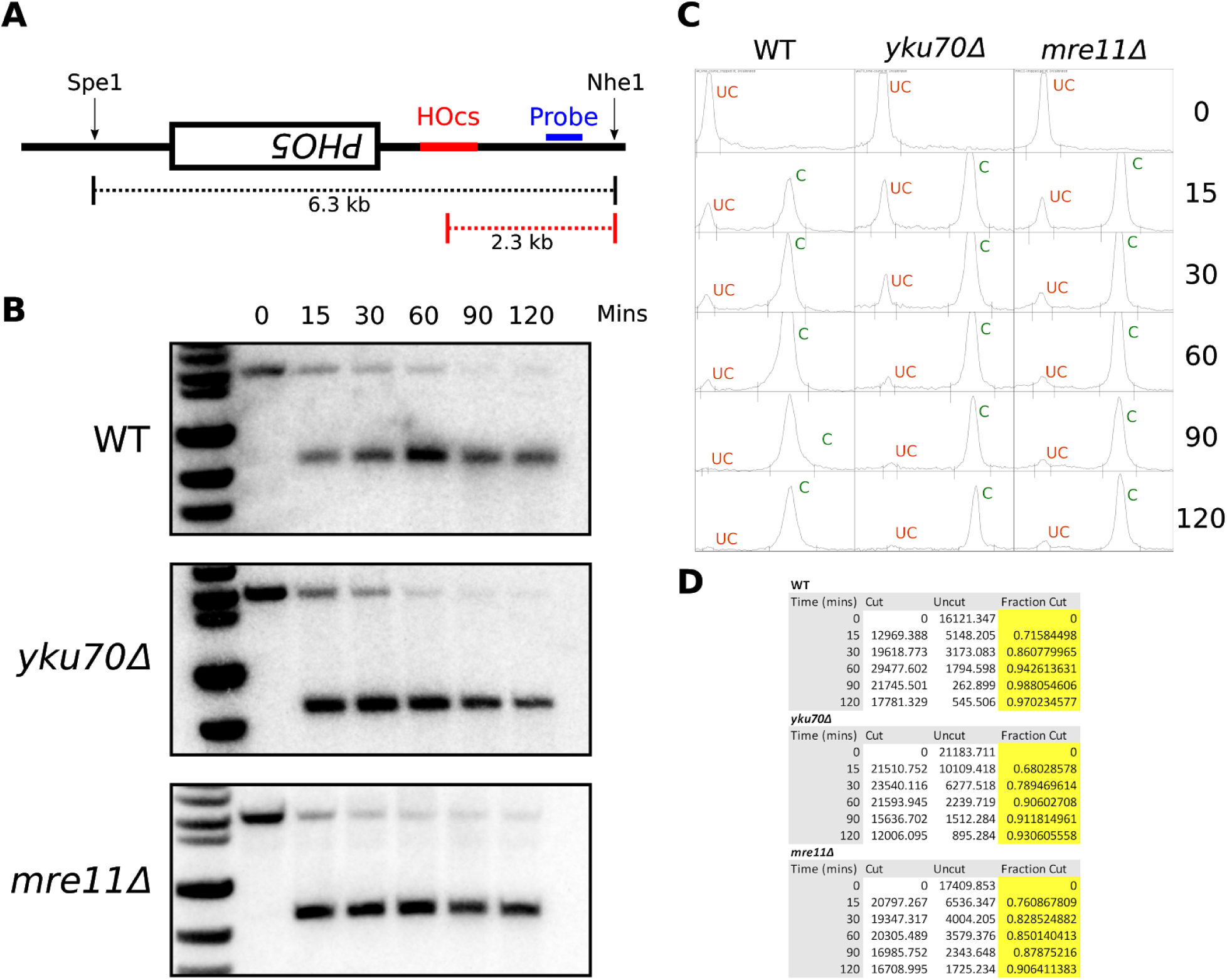
Southern Blotting of WT, yku70*Δ*, and mre11*Δ* Strains. **A.** Schematic of the southern blotting probe and the detected fragment sizes at the *PHO5* locus with the 117 bp HO recognition site. **B.** Southern blots of the 3 strains profiled in this study. **C.** Line profiles of each lane that were analyzed in ImageJ to extrapolate uncut (UC, orange) and cut (C, green) band intensity. **D.** Table of the data obtained from the analysis in **C**, fraction cut is determined by dividing the cut signal by the sum of the uncut and cut signals.

**Supplemental Figure 3.**
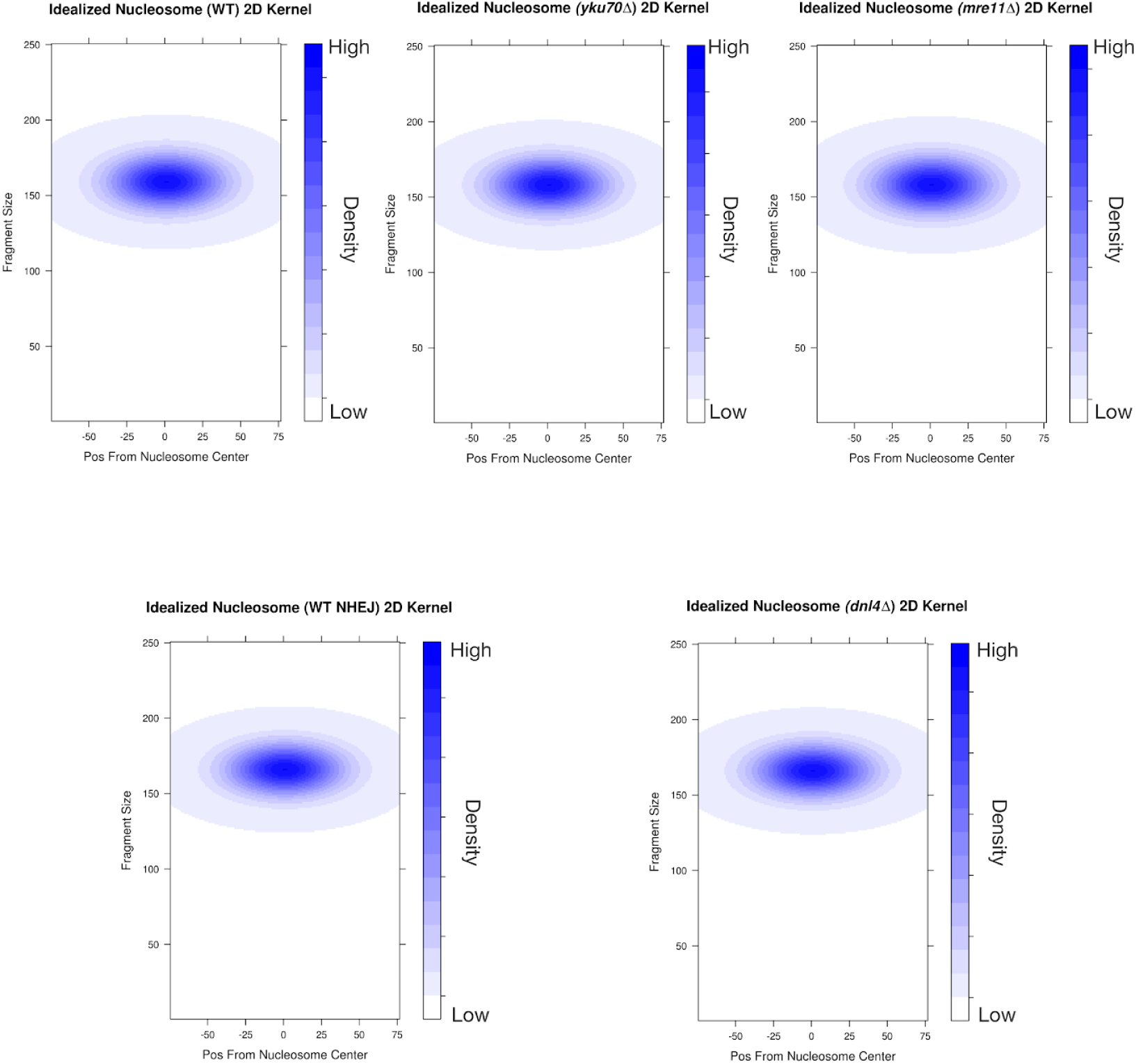
Idealized two-dimensional density (2D) kernel of nucleosomes derived from the pre-induction sample for each time-course. The size and fragment distribution parameters for this nucleosome were derived from analyzing the 8,632 nucleosomes chemically mapped by Brogaard, et al.(Brogaard et al. 2012) on chr IV for each experiment and were similar across all experiments.

**Supplemental Figure 4.**
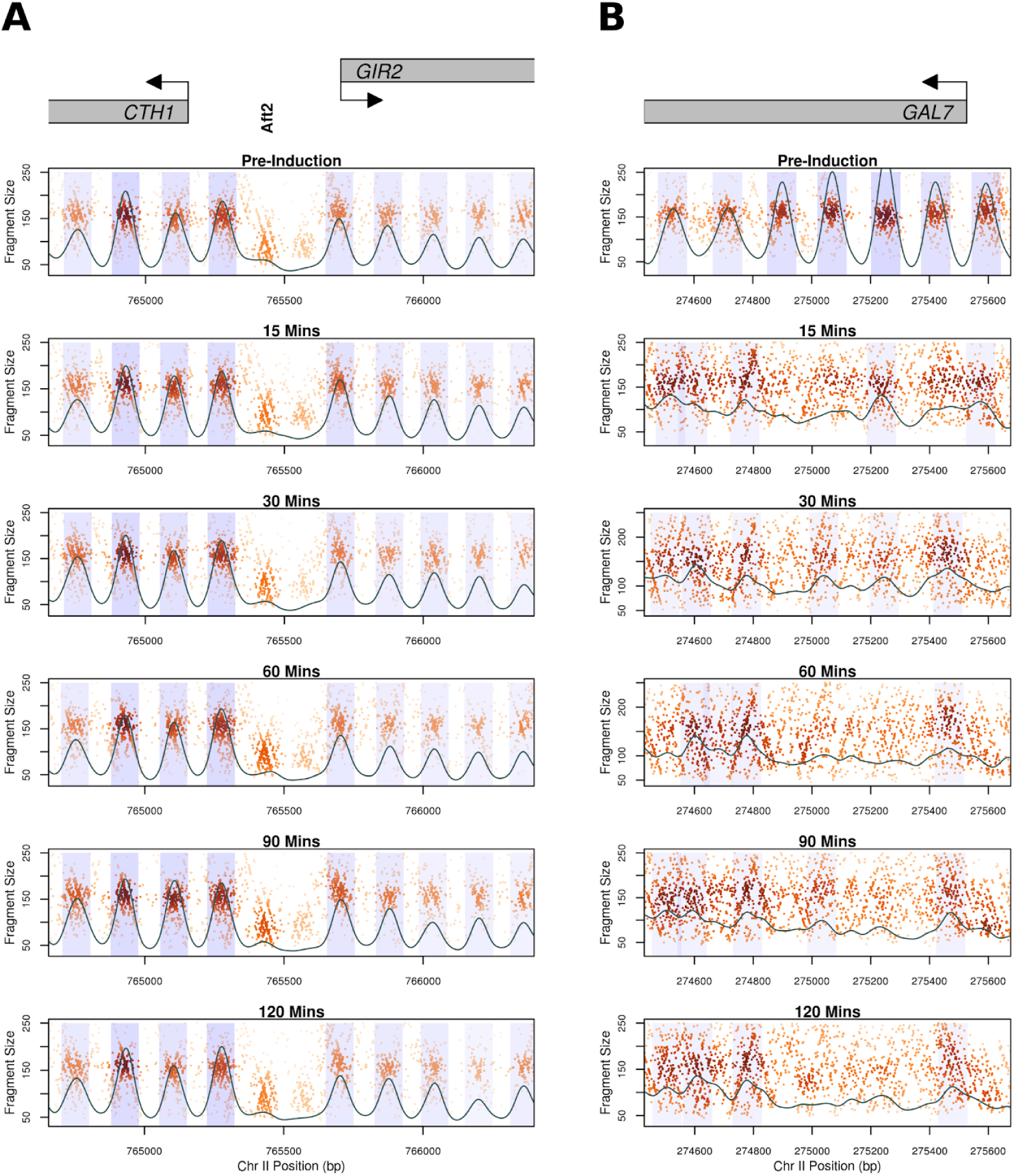
*yku70Δ* Control Loci Genome Wide Chromatin Occupancy Profiles (GCOPs). **A.)** GCOP for the *CTH1* and *GIR2* loci which serve as a control for unaltered chromatin. Gray boxes depict gene bodies, with arrows indicating the direction of transcription. The bold vertical text indicates the Aft2p binding site in between these two genes. A two-dimensional cross correlation with an idealized nucleosome (see methods) is calculated at every base pair and depicted as a continuous black trace. The peaks of this cross-correlation analysis represent the most likely position of a locally detected nucleosome and we have shaded +/− 0.5 standard deviation of nucleosome position from the center of each detected peak. The intensity of the shaded color is proportional to the nucleosome’s peak cross-correlation score. **B.** GCOP for the *GAL7* locus following galactose induction.

**Supplemental Figure 5.**
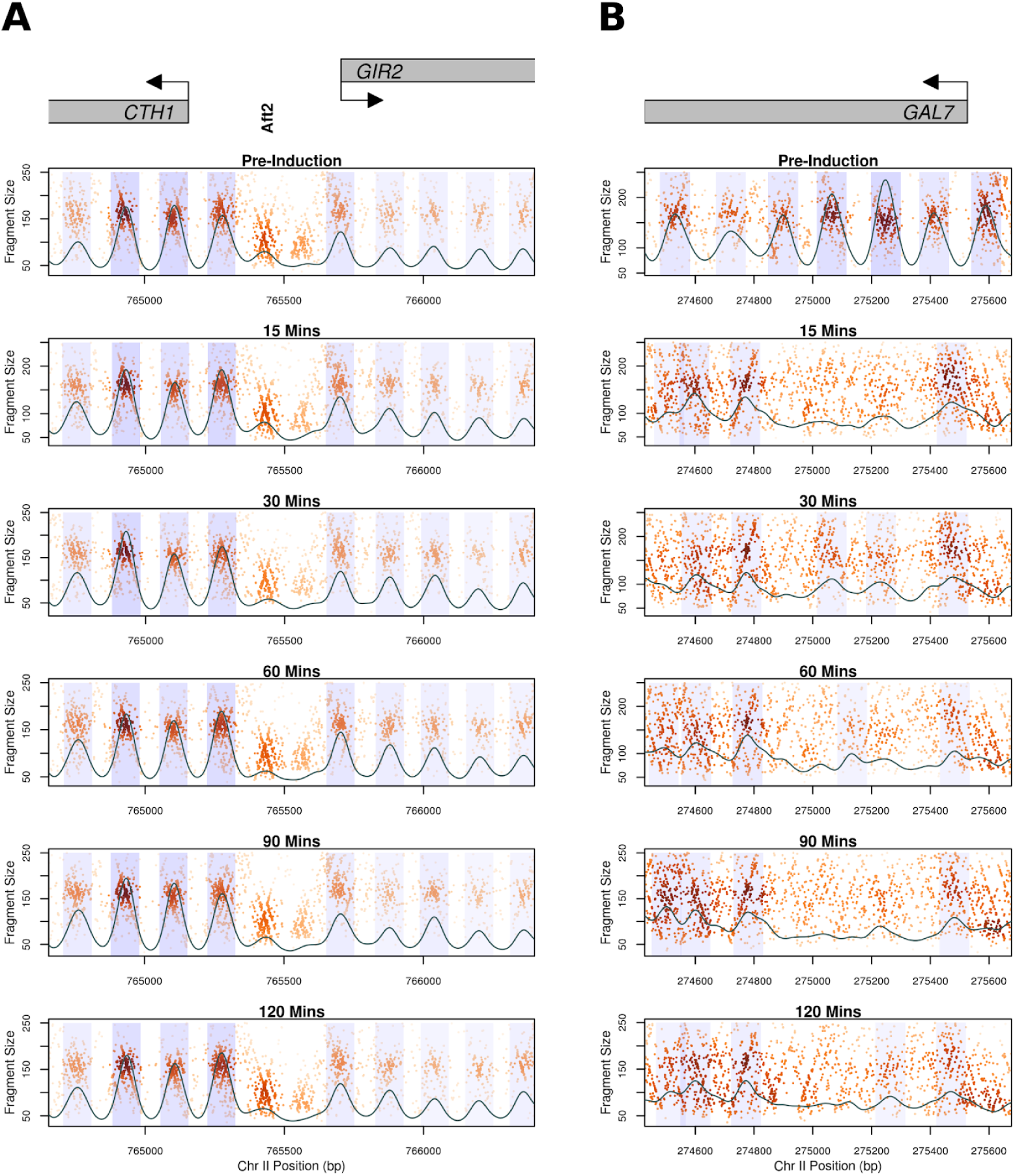
*mre11Δ* Control Loci Genome Wide Chromatin Occupancy Plots (GCOPs) **A.)** GCOP for the *CTH1* and *GIR2* loci which serve as a control for unaltered chromatin. Gray boxes depict gene bodies, with arrows indicating the direction of the ORF. The bold vertical text annotatesthe Aft2p binding site in between these two genes. A two-dimensional cross correlation with an idealized nucleosome (see Methods) is calculated at every base pair and depicted as a continuous black trace. The peaks of this cross-correlation analysis represent the most likely position of a locally detected nucleosome and we have shaded +/− 0.5 standard deviation of nucleosome position from the center of each detected peak. The intensity of the shaded color is proportional to the nucleosome’s peak cross-correlation score. **B.** GCOP for the *GAL7* locus following galactose induction.

**Supplemental Figure 6.**
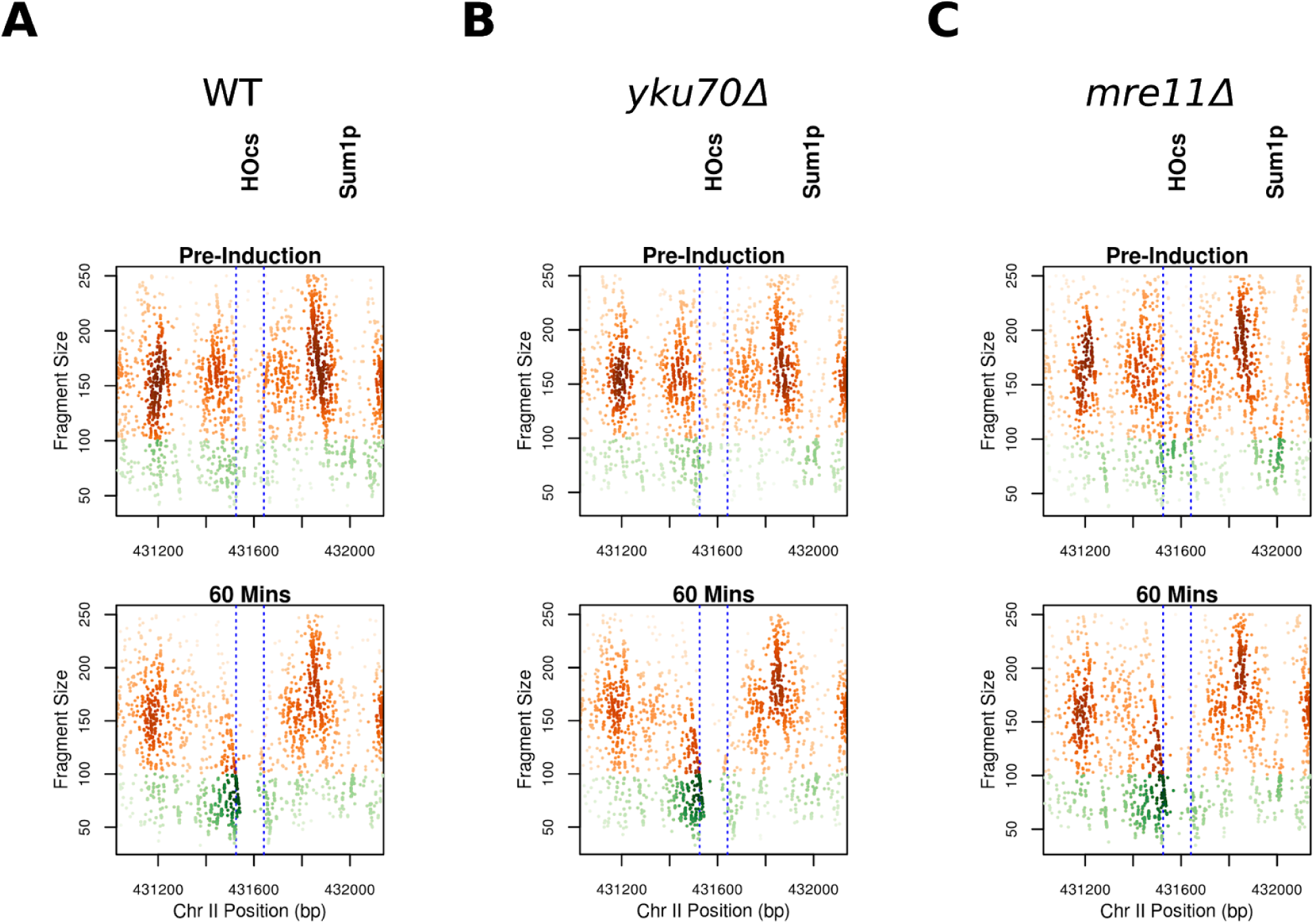
Zoomed-in view of genome wide chromatin occupancy plots of the WT (**A**), *yku70Δ* (**B**), and *mre11Δ* (**C**) strains. Fragments smaller than 100bp (sub-nucleosomal) are colored in green.

**Supplemental Figure 7.**
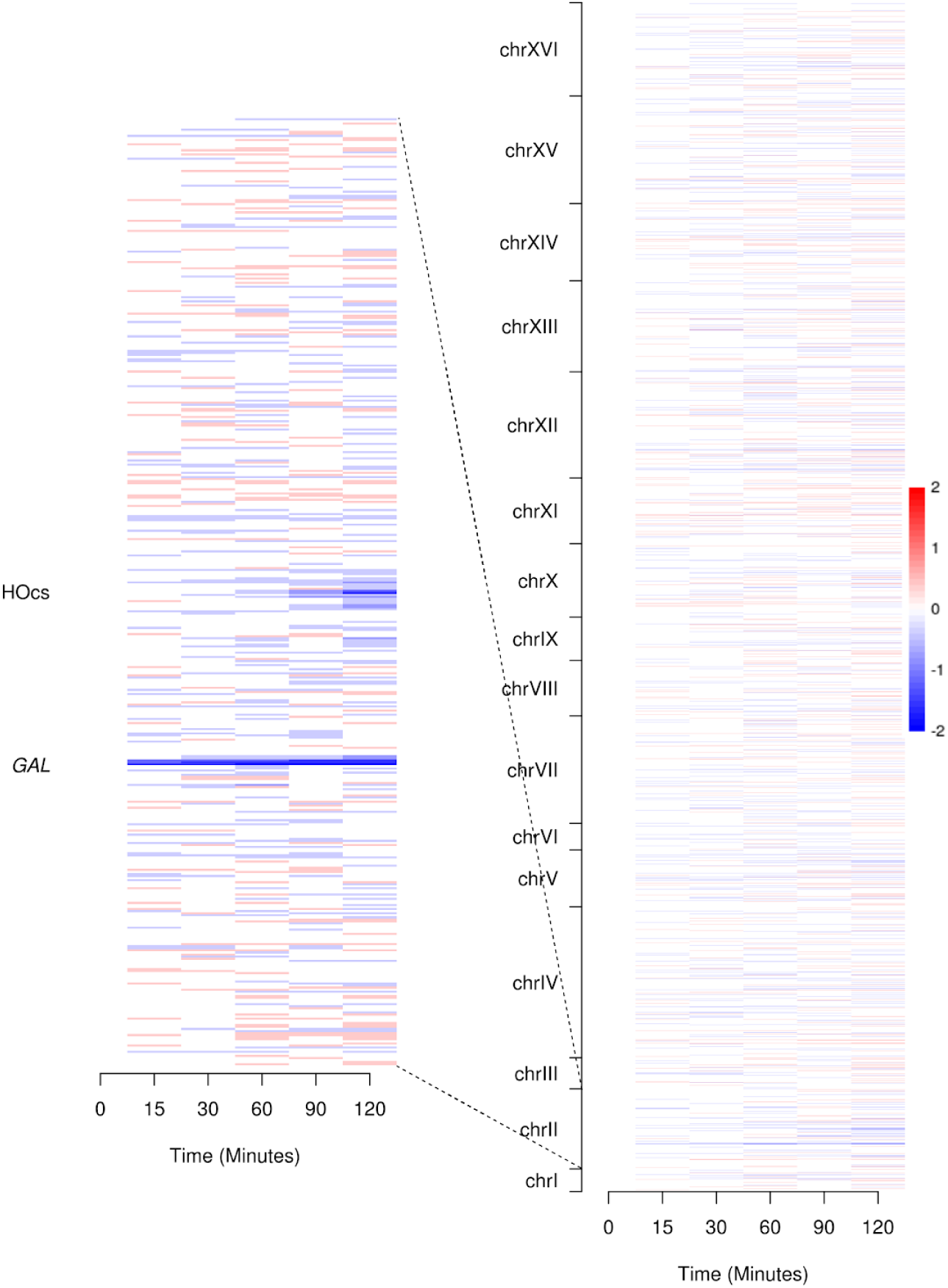
Genome wide genic nucleosome analysis in wild type cells following a DSB. The 2D cross correlation score (similarity to an idealized nucleosome) for genic nucleosomes in each time point is expressed as a log2-fold change relative to the pre-induction state. The coordinates corresponding to the *GAL* loci and the Ho induced DSB are annotated on the zoomed in view of Chr II (left panel).

**Supplemental Figure 8.**
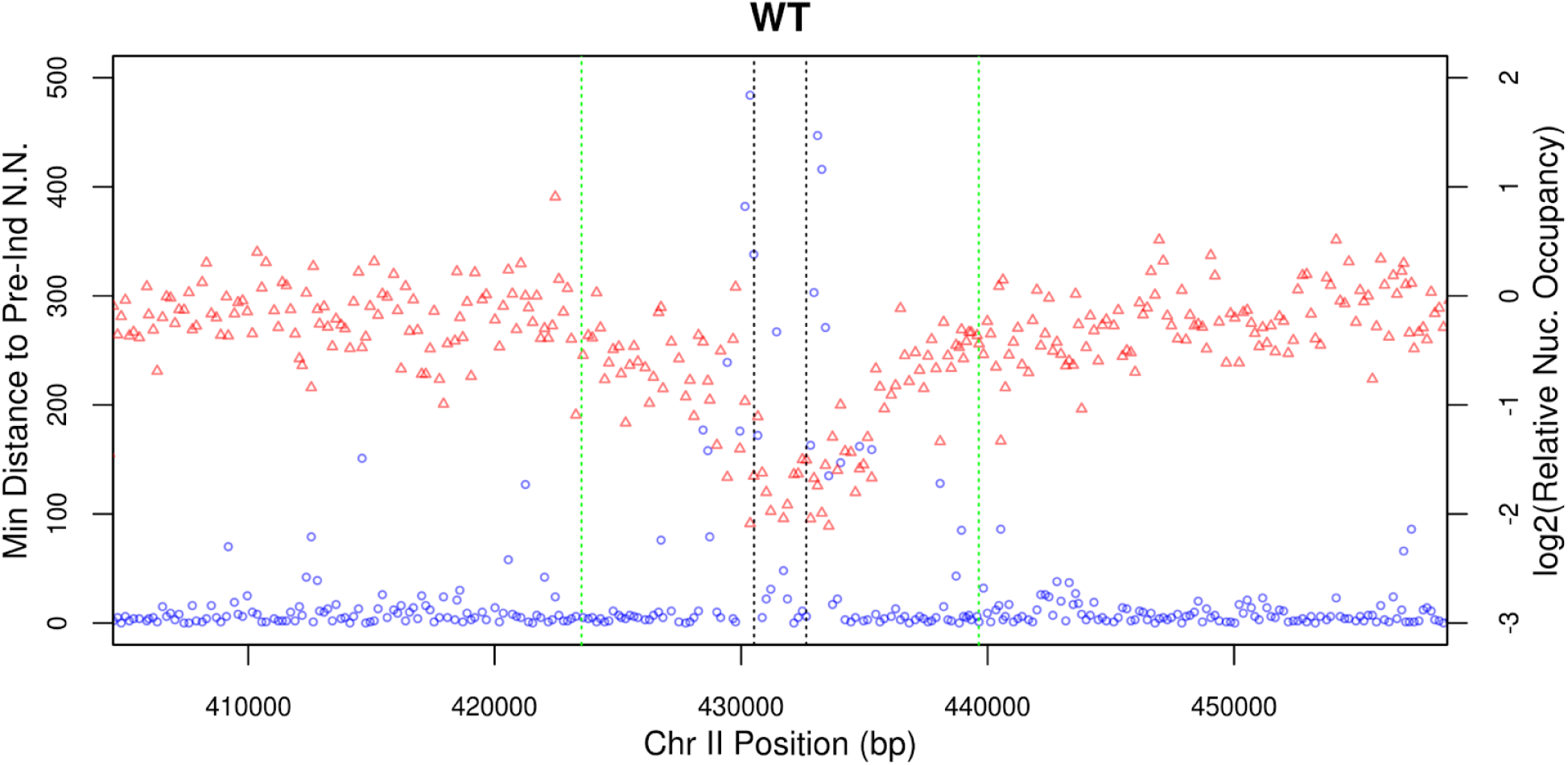
Relative nucleosome position and nucleosomal occupancy ∼25kb from the DSB. We detected ∼4700 nucleosomes in an ∼50kb window around the DSB (arrayed on the x axis according to their position) on Chr II and plotted their relative distance to the nearest neighbor nucleosome (in pre-induction) throughout the time course (blue circles). Separately we calculated the occupancy of each individual nucleosome and plotted the log2-ratio of the 120 min post-induction occupancy to the pre-induction occupancy (red triangles). The dotted green lines denote coordinates 8kb from the break while the dotted black lines denote 1kb from the break.

**Supplemental Figure 9.**
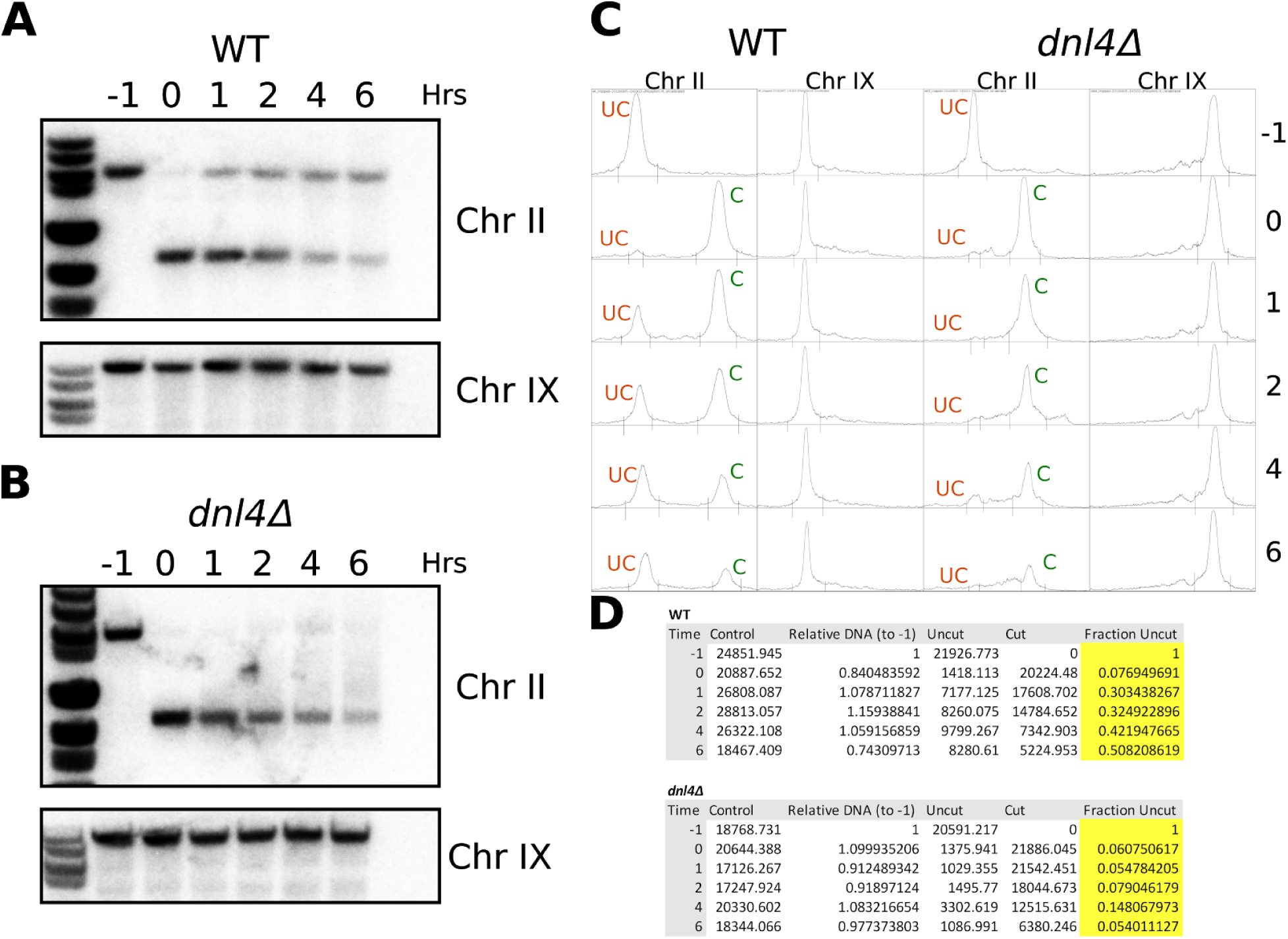
Southern Blotting of WT and *dnl4Δ* strains. Southern blots of the WT (A) and *dnl4Δ* (B) strains along with an unrelated control locus from Chr IX. **C.** Line profiles of each lane that were analyzed in ImageJ to extrapolate uncut (UC, orange), cut (C, green) and control (Chr IX) band intensity. **D.** Table of the data obtained from the analysis in **C**, fraction uncut is determined by dividing the uncut signal at each time point by the uncut signal from pre-induction. All of these values are then normalized to the band intensity from Chr IX in the corresponding lane.

**Supplemental Figure 10.**
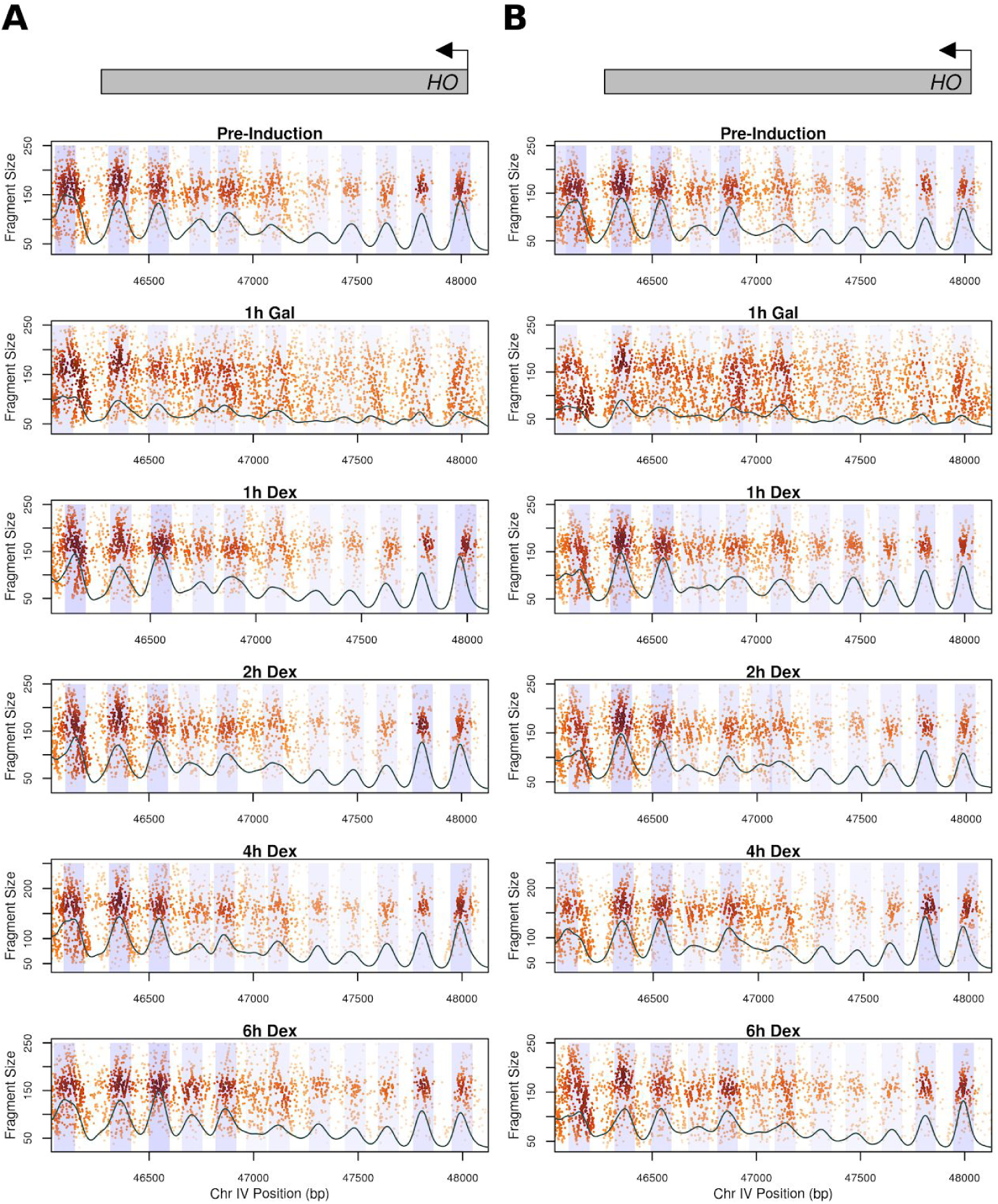
Genome Wide Chromatin Occupancy Plots (GCOPs) of the HO gene body in the WT (**A**) and *dnl4Δ* (**B**) strains over the NHEJ time course. Gray boxes depict gene bodies, with arrows indicating the direction of transcription. A two-dimensional cross correlation with an idealized nucleosome (see Methods) is calculated at every base pair and depicted as a continuous black trace. The peaks of this cross-correlation analysis represent the most likely position of a locally detected nucleosome and we have shaded +/− 0.5 standard deviation of nucleosome position from the center of each detected peak. The intensity of the shaded color is proportional to the nucleosome’s peak cross-correlation score.

**Supplemental Figure 11.**
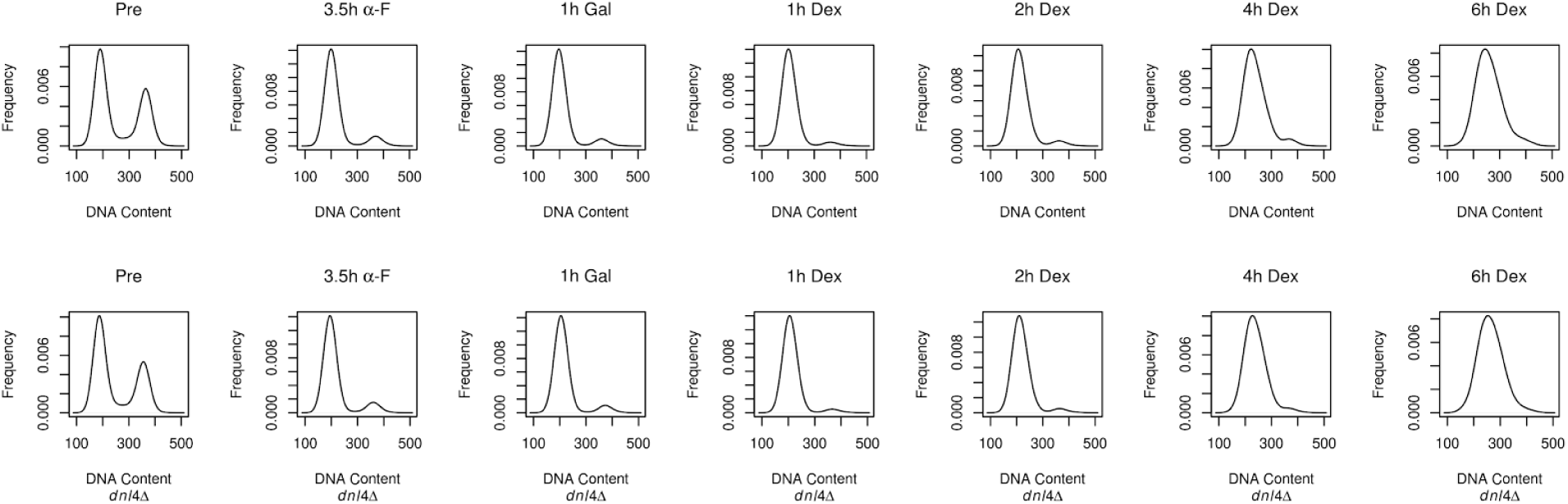
FACS profiles of the WT (top) and *dnl4Δ* (bottom) strains throughout the NHEJ time course. DNA content is measured on the horizontal axis while frequency is measured on the vertical axis.

